# A complete account of the behavioral repertoire uncovers principles of larval zebrafish hunting behavior

**DOI:** 10.1101/2024.10.05.613699

**Authors:** Yoav Rubinstein, Maayan Moshkovitz, Itay Ottenheimer, Sapir Shapira, Stas Tiomkin, Lilach Avitan

## Abstract

In goal-directed behavior animals select actions from a diverse repertoire of possible movements. Accurately quantifying the complete behavioral repertoire can uncover the underlying rules that guide such goal-directed behavior. However, these movements are usually complex, high-dimensional, and lead to various outcomes, posing a challenge to fully capture the complete repertoire. By tracking freely hunting zebrafish larvae using a highspeed camera and analyzing their movements, we developed a mathematical model that accurately reproduces the complete repertoire. Using the model, we show that fish position and change in heading angle following a movement are coupled, such that the choice of one of them limits the possibilities of the other. This repertoire structure uncovered fundamental principles of movements, showing that fish rotate around an identified rotation point and then move forward or backward along straight lines. From the uncovered movement principles, we identified a new guiding rule for prey interaction: in each movement, fish turn to face the prey and then move forward or backward. This enables decoupling between orientation and distance selections of the fish during the hunt. These results provide a comprehensive and continuous description of the repertoire of movements, reveal underlying algorithmic rules that govern the behavior, and offer insights into the potential neural implementation of the repertoire.

## Introduction

Natural goal-directed behavior involves executing a sequence of actions to achieve a specific goal. By acting, the animal progressively improves its relation to the goal until it has been achieved. The actions available to the animal constitute its behavioral repertoire. The behavioral algorithm, or the rules the animals follow to interact with the goal, determines which action is selected from this repertoire given the goal. To uncover the behavioral algorithm implemented by the animal, it is crucial to thoroughly characterize the range of possible actions available to the animal.

Actions are often studied from two complementary perspectives : (i) quantifying the dynamics of recorded motor movements, i.e., movement dynamics of the paws, wings, limbs, or tail ^2,4,12,24,31,40^, and (ii) quantifying the outcome of each movement, i.e the impact of a movement on the state of the animal in the world ^2^. While movement dynamics are often high-dimensional and complex, their outcomes are usually lower-dimensional and easier to study ^7,37^. Therefore, outcomes are more amenable to being captured using a straightforward mathematical rule. This mathematical description can account for the available repertoire of the animal and form the basis for unraveling its behavioral algorithm.

The natural larval zebrafish hunting behavior is a powerful model for studying the behavioral repertoire and the underlying behavioral algorithm. A hunting event proceeds via a sequence of discrete movements, thus single movements are easily segmented. With each movement, larvae refine prey localization in their visual field until they capture the prey ^8,9,15,16,26,37^. Understanding the behavioral rule that larvae use to achieve this refinement has been the focus of several works^7,15,21,27,32,37^. While it is clear that the animal aims to capture the prey and closely interacts with it, our ability to predict the fish selection of position and orientation in each action is still limited.

Most approaches to studying fish movements have focused on tail movements ^8,9,10,17,21,22,24,25,27,28,30,32^, although fins and jaw movements have also been studied ^13,19,25,27,36^. These tail movements form a high-dimensional continuum in the tail features space. Therefore, movements were clustered into several discrete types ^17,21,24,27^. These techniques have provided insights into the possible types of movements, the available repertoire, and the process of chaining movements from different clusters ^17,21,24,27^. However, determining where the fish is in space (position and orientation) following a movement of a particular type is currently impossible. As a result, our understanding of the behavioral algorithm as reflected by the prediction of a specific movement selection during the hunt remains limited.

An alternative to describing the behavioral repertoire entails studying it from an outcome perspective ^7,18,37^. Rather than focusing on the movement of the tail, eyes, fins and jaw, all of which contribute to the change in position and orientation, this approach focuses on the possible positions and orientations the fish can achieve after a single movement. Therefore, it provides a continuous and low-dimensional description of the repertoire. The possible combinations of position and orientation parameters define the repertoire of all possible outcomes. However, so far outcome parameters have been studied as independent parameters ^7,18,37^. Therefore, a comprehensive understanding of these parameters, their relation, and the repertoire of outcomes they form are still missing. For instance, for a given change in heading angle - a well-known outcome parameter with an identified neural correlate ^33^ - it is unclear where in space the fish can be positioned.

The dimensionality of the behavioral repertoire is crucial to understanding the repertoire itself ^11,35^, the process of action selection, and the underlying visuomotor transformation. Recently, hindbrain neural activity related to eyes and tail movements in response to a rotating striped pattern was found to be two-dimensional ^14^. While this suggests that the repertoire of movements elicited can potentially be low-dimensional, it remains unclear whether this finding can be extended to the complete natural repertoire. Measuring the dimensionality of the movement repertoire in natural settings is challenging due to the high-dimensionality of tail movements ^17,24,27^. While focusing on outcomes offers the potential to quantify the repertoire in a more simplified, low-dimensional manner, this approach has remained unexplored.

A complete and closed mathematical description of the repertoire can aid in uncovering the underlying behavioral algorithm that fish implement to capture the prey. When interacting with the prey, larvae were reported to turn at an angle that is about 40%-60% of the prey angle in each movement ^7,37^. While this rule defines the change in orientation, fish position can vary for a given change in orientation. Therefore, for a given prey angle and a respective (60%) change in fish orientation ^7,37^, some fish positions will be better than others with respect to the prey. This suggests that orientation selection by the fish may impact position selection. Therefore, understanding the relationship between the change in orientation and position following a movement is crucial to understanding the behavioral algorithm employed by hunting larvae.

Here, we show that three variables are essential to fully describe the position and orientation of the fish following a movement. We provide a mathematical model that accurately produces the entire outcome repertoire and uncovers the relation between outcome parameters in natural hunting behavior settings. We show that fish position following a given movement predicts the associated change in heading angle. In addition, we demonstrate that a given change in heading angle dictates a limited set of possible positions. This coupling between position and orientation suggests that outcomes are two-dimensional. By linking outcomes to movements, we show that fish rotate around an identified rotation point and then move forward or backward along a straight line. This nature of movements creates the coupling between orientation and position. Based on the uncovered repertoire, we show that for most hunting movements, fish close the entire angle to the prey when this prey angle is measured relative to the uncovered rotation point. We therefore suggest a new explanation for how fish select a change in heading angle, proposing that they aim to face their prey, which enables the independent selections of orientation and position.

## Results

### Bout azimuthal angle is crucial to describe fish position and orientation following a movement

To uncover the behavioral repertoire available to the larval zebrafish during hunting, we recorded natural hunting behavior for 15 minutes (18 fish, 5-7 days post fertilization) using a high-speed camera^3^ (See Methods). We utilized our custom-developed tracking system and extracted various features from each frame (See Methods). This included the contours of the fish, eyes, heading direction, swim bladder, and tail midline, as well as features related to the prey in the dish, such as their positions, trajectories, and identification of the target prey during each hunting event (Figure 1A, Supplementary Figure S1A, Supplementary Movie1, Movie2). In addition, the discrete nature of larval zebrafish movement ^39^ enabled the segmentation of individual movements along the hunting event (See Methods). The tracking system extracted 4006 movements across 939 detected hunting events (See Methods).

**Figure 1:**
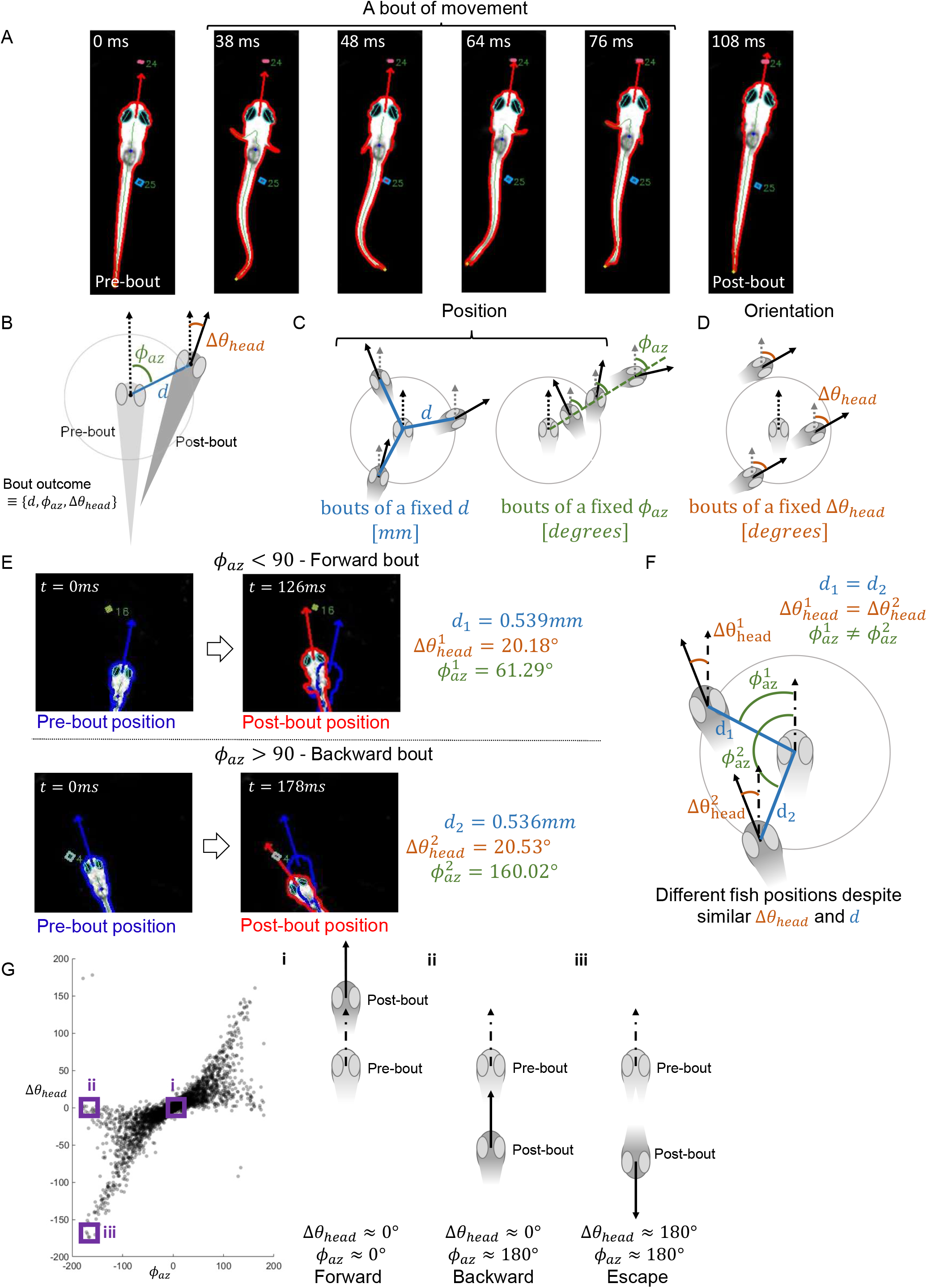
Bout outcomes are defined by three parameters. **A:** A set of six frames from a single fish movement during a hunting event. The coordinated movements of the tail, fins and eyes lead to a specific change in position and orientation (an outcome). Fish contour, head direction, fitted eye ellipses, swim bladder, tail midline, and tail tip were tracked and indicated on the frames. All prey items were detected and tracked, and two such prey items were indicated on the frames. **B:** A bout outcome is described by three parameters: bout distance (*d*, blue line), the change in heading direction following the bout (Δ*θ*_*head*_, orange angle), and azimuthal angle (*ϕ*_*az*_, green angle). In all panels throughout the figure pre- and post-bout fish poses (fish orientation and position) are indicated by a bright and a dark grey fish heads respectively. Pre- and post-bout heading directions are marked with dashed and solid arrows, respectively. **C:** Left: example fish poses following a bout of a fixed distance (*d*) and varying changes in heading angle (Δ*θ*_*head*_) and azimuthal angles (*ϕ*_*az*_). Post-bout poses are positioned on the perimeter of a circle. Distance throughout the paper is measured in mm. Right: example fish poses following a bout of a fixed azimuthal angle (*ϕ*_*az*_) with different distances (*d*) and different changes in heading angle (Δ*θ*_*head*_). Post-bout poses are positioned on the green dashed line which determines a particular azimuthal angle (green angles). Azimuthal angle throughout the paper is measured in degrees. **D:** Example fish poses following a bout of a fixed change in heading angle (Δ*θ*_*head*_) with different bout distances (*d*) and azimuthal angles (*ϕ*_*az*_). The change in heading angle throughout the paper is measured in degrees. **E:** Two example bouts (top and bottom) with pre-bout and post-bout poses marked by the blue and red contours respectively. Both bouts were of similar changes in heading angle and distance. However, the fish moved forward in the top example and backward in the bottom example as captured by the different azimuthal angles. **F:** A schematic of the two example bouts showing that despite similar changes in heading angle and bout distance, fish can be placed in different positions in space due to the different bout azimuthal angles. **G:** The relation between the change in heading angle and the azimuthal angle for all bouts in our dataset shows that these angles can have different values relative to one another. Three example combinations of these two angles marked in three purple boxes represent forward (Gi), backward (Gii) and escape (Giii) bouts. Actual movements representing the three examples are shown in Supplementary Movie 4.

Every bout of movement recruited coordinated tail, fins, eyes, and occasionally jaw movements (Figure 1A). Despite the complexity of these movements, each movement resulted in a specific simple change in the position and orientation of the fish in space. We termed this change as the movement outcome. We quantified movement outcomes using three parameters in polar coordinates: the change in heading angle (Δ*θ*_*head*_), the distance traveled (*d*), and the azimuthal angle (*ϕ*_*az*_), defined by the angle between the pre-bout heading direction and the movement direction (Figure 1B) (See Methods). The pair *d* and *ϕ*_*az*_ describe fish position following a movement (Figure 1C), and Δ*θ*_*head*_ accounts for the change in orientation relative to the pre-bout heading angle (Figure 1D). Since the environment in this experiment is approximately two-dimensional (the chamber is 25 mm deep and was imaged from the top), using these three parameters (*d, ϕ*_*az*_, and Δ*θ*_*head*_) we can fully reconstruct movement outcomes (position and orientation of the fish) following any given movement.

These three parameters were previously studied both in the context of movement characterization and in relation to prey features ^7,18,37^. Since movement distance (*d*) and the change in heading angle (Δ*θ*_*head*_) were the most frequently studied parameters ^18,37^, we asked whether we could reconstruct the full outcome of any movement using these two parameters. More specifically, we sought to determine whether we could resolve the exact position and orientation of the fish following any movement using bout distance and the change in heading angle. We found that following movements of a similar change in heading angle and distance, fish can be positioned in different positions in space due to distinct changes in azimuthal angle (Figure 1E,F, Supplementary Movie 3). Therefore, the azimuthal angle is crucial for accurately reconstructing movement outcomes.

This azimuthal angle differs from the change in heading angle (Figure 1G), and different combinations of these angles lead to qualitatively distinct outcomes. For instance, forward movements are characterized by no change in fish orientation and movement in the pre-bout heading direction, such that the change in heading angle and the azimuthal angle are near zero (Figure 1Gi, Supplementary Movie 4). Backward movements are characterized by no change in fish orientation with the fish moving in the opposite direction of the heading direction. Therefore, the change in heading angle is near zero and the azimuthal angle is close to 180°(Figure 1Gii, Supplementary Movie 4). In escape movements, which occasionally occur at the end of the hunting event (47% of hunting events), fish orient and move away from their original heading direction, resulting in both angles being near 180°(Figure 1Giii, supplementary Movie 4). Together, the azimuthal angle contains information not captured by the change in heading angle and distance, making it essential for fully reconstructing the outcome of a given movement.

### A change in fish position dictates a particular change in heading angle

While the change in heading angle, the azimuthal angle, and the distance traveled following a movement were studied as independent variables^7,37^, they are a consequence of a single fish movement. Therefore, their relation is crucial to uncovering the repertoire of possible outcomes. To uncover the relation between the three parameters, we aligned detected bouts during all hunting events according to the pre-bout pose (Figure 2A, middle black dot for position and dashed black arrow for orientation). We then examined all possible bout outcomes relative to this pre-bout pose (Figure 2A, dots represent the different positions, colored arrows represent the change in heading angle). While fish moved to a range of positions (represented by the dots and defined by *ϕ*_*az*_ and *d*), neighboring fish positions (nearby dots) showed a similar change in heading angle (similar colors), suggesting a correlation between the change in position and the change in heading angle. This correlation indicated that not all outcomes (or combinations of the three parameters) are possible following a movement.

**Figure 2:**
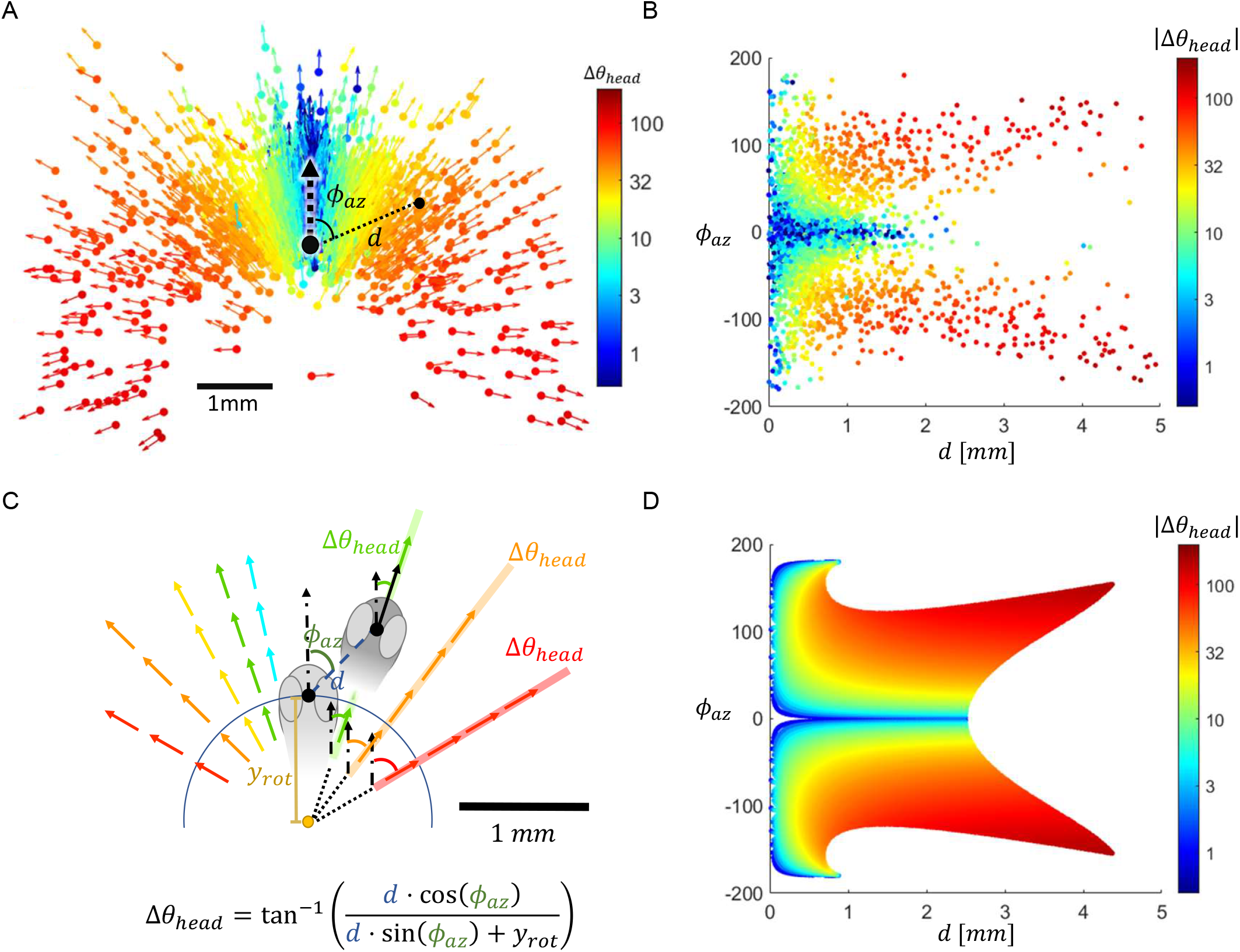
A simple mathematical model captures the repertoire of bout outcomes. **A:** The measured repertoire of bout outcomes. Pre-bout positions and orientations were aligned such that they are indicated by the black dot and the dashed black arrow respectively. The outcome of each bout is represented by a colored dot and arrow. The dots represent the post-bout position (determined by bout distance and azimuthal angle relative to the pre-bout position and heading direction), and the direction of arrows represents the change in heading angle. Arrows and dots are colored according to the change in heading angle, showing that neighboring post-bout positions (close-by dots) share a similar change in heading angle. The absolute change in heading angle is colored on a log scale. **B:** A representation of the outcome repertoire. Each movement is represented by a single dot specifying the post-bout position (by *ϕ*_*az*_ and *d*). The dots are colored by the change in heading angle. Colors scheme is similar to Figure 2A. For a specific change in heading angle (a specific color), there is a limited set of positions defined by the pair azimuth and distance. **C:** Top: a mathematical model of the outcome repertoire uncovers the relation between position and orientation parameters. Fish positions following movements of a similar change in heading angle, indicated by the same color, are aligned along a single straight line. Lines representing different changes in heading angle converge at a single point, positioned at a distance *y*_*rot*_ behind the midpoint between the eyes. Bottom: the change in heading angle is calculated from the positional parameters using trigonometric relations. **D:** The model prediction of change in heading angle given the positional parameters reconstructs the empiric relations between position and orientation (compare to panel 2B). Error quantification of the model and a 3D structure of the repertoire are shown in Figure S2A-D and Supplementary Movie 5.

The relation between the positional parameters (*d, ϕ*_*az*_) and the change in heading angle (Δ*θ*_*head*_) can be further observed when outcomes are presented according to their positional parameters (*d, ϕ*_*az*_) and colored according to the change in heading angle (Figure 2B). This visualization emphasizes that for a given change in heading angle (a particular color), larvae have a limited set of positional options defined by the pair distance and azimuthal angle. This further suggests that the positional parameters (*d, ϕ*_*az*_) can potentially reconstruct the full outcome, as the change in heading angle may be derived from the change in position.

To quantitatively capture the relation between fish position and change in heading angle following a movement, we used two observations from the repertoire of possible outcomes following a bout (shown in Figure 2A). The first observation suggests that all possible positions of the fish for a given change in heading angle may align along a single linear line (this linear line is observed for small changes in heading angle and is less prominent for larger angles due to the logarithmic color scale). Thus, every change in heading angle establishes a different linear line of possible positions. The second observation suggests that the extrapolations of these linear lines meet at a single point. This point was found behind the midpoint between the eyes at a distance which we termed *y*_*rot*_ (Figure 2C). Based on these observations, we identified a model with a single free parameter (*y*_*rot*_) that fits all outcomes in our data (Figure 2C, bottom). The fitting process, which was applied to all fish together, yielded a value of *y*_*rot*_ = 1 ± 0.03*mm* (See Methods). This model thus offers a closed mathematical relation that predicts the change in heading angle given the post-bout position of any movement.

The model accurately reconstructed the repertoire of measured outcomes (Figure 2D, compared to Figure 2B). Specifically, for each post-bout position, the model was able to accurately predict the change in heading angle with a prediction error of less than 4 degrees in 76% of the bouts (Figure S2A). Additionally, for a given change in heading angle, the model predicted a set of post-bout positions arranged along a linear line. Empiric post-bout positions, for any change in heading angle, were placed up to 0.2 mm from the predicted lines in 91% of the bouts (Figure S2B). Together, the model captures the relation between positional parameters (*d, ϕ*_*az*_) and the change in heading angle (Δ*θ*_*head*_). It demonstrates that the change in heading angle can be accurately derived from the positional parameters. Furthermore, it shows that the change in heading angle determines a set of possible fish positions aligned along a specific linear line that we can accurately define.

Our model uncovered that all possible outcomes lie within a 2D manifold in the 3D outcome parameter space (Figure S2C, Supplementary Movie 5). More specifically, the theoretical relation between the parameters (Figure 2C) suggested that all outcomes were determined by two parameters: distance and azimuthal angle. Interestingly, this relation also indicated that other combinations of two parameters (distance and change in heading angle or change in heading angle and azimuthal angle) would be insufficient to reconstruct the outcomes of all movements (as shown in Figure S3). We quantified how well we can reconstruct the full outcome using the other two parameters in our data. The change in heading angle and distance could not resolve a unique azimuthal angle for 16% of the bouts (Figure S3E), while the change in heading angle and azimuthal angle could not resolve a unique distance for 13% of the bouts (Figure S3F). Therefore, outcomes are fully and uniquely described using the positional parameters (*ϕ*_*az*_ and *d*), while other pairs of parameters can only partially capture the full repertoire.

### The outcome repertoire reveals principles of movements

Next, we aimed to interpret the uncovered repertoire in terms of movements. The model relied on the point at a distance *y*_*rot*_ behind the eyes, where all lines representing different changes in heading angle met. We assumed that this point was the rotation point of the fish. Under this assumption, movements can be conceptualized as the fish rotating around this rotation point and then moving forward or backward along a line determined by the change in heading angle (Figure 3A). We asked whether the actual movement of the fish would agree with this interpretation. We collected bouts of different changes in heading angle and examined the trajectories of the midpoint between the eyes along the time course of the movement. The rotation of the fish around the rotation point was manifested by the trajectory of the point between the eyes drawing circular arcs centered at *y*_*rot*_, with longer arcs for larger changes in heading angle (Figure 3B). Each arc terminated on the linear line predicted by the change in heading angle. Therefore, the circular phase of the trajectory confirmed that the larvae rotate around the hypothesized rotation point.

**Figure 3:**
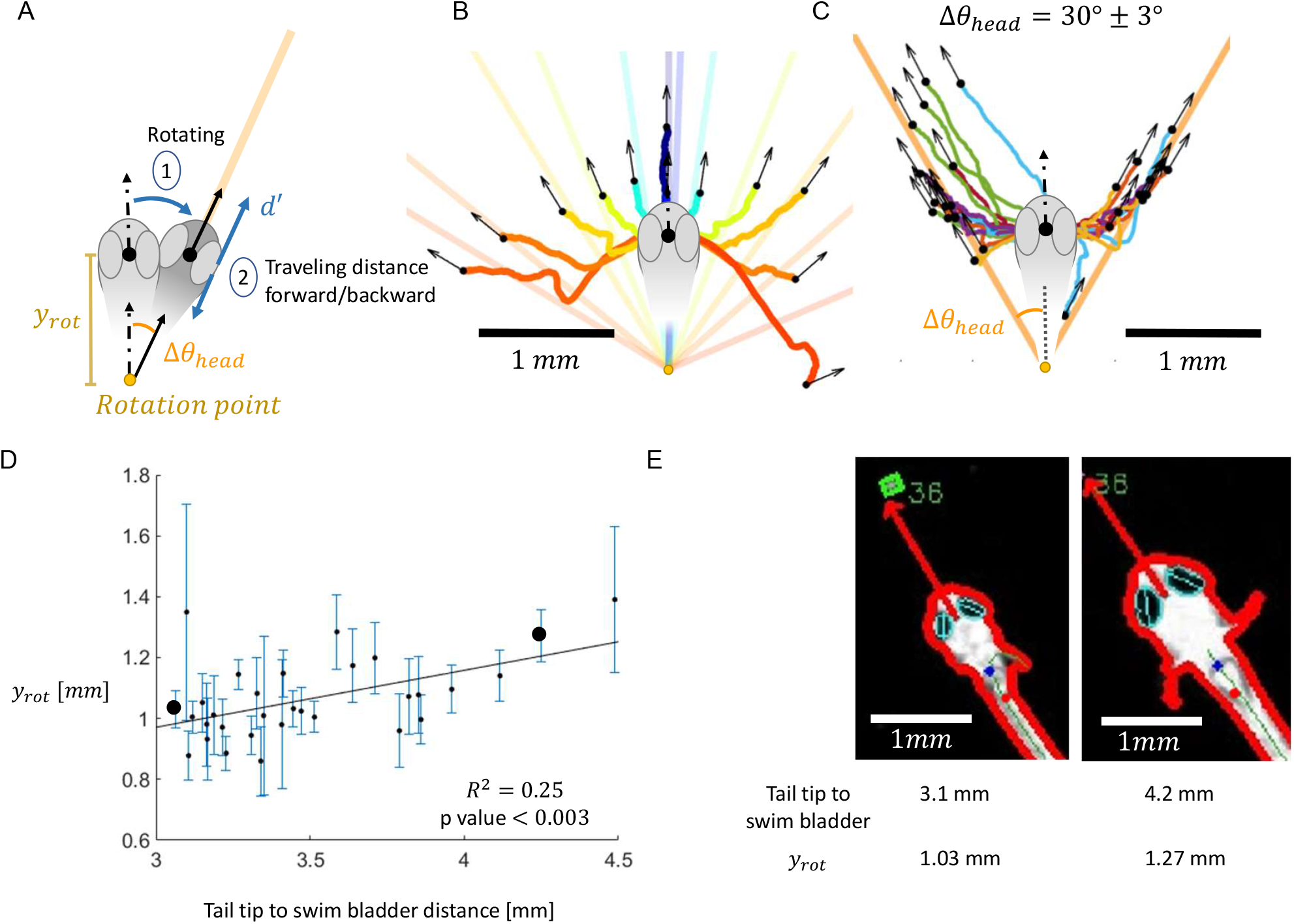
The repertoire of movement outcomes uncovered principles of movement trajectories. **A:** A possible interpretation of the modeled repertoire. The movement has two main phases, including rotation and movement forward or backward. In the first phase, fish rotate (Δ*θ*_*head*_) around a rotation point found at a distance *y*_*rot*_ behind the midpoint between the eyes. The rotation specifies a line of possible positions (orange line). In the second phase, fish move a distance *d*^*′*^ along the line specified by the rotation angle (orange line). This results in a set of possible fish positions along a certain line defined by a specific change in heading angle. **B:** The trajectory of the midpoint between the eyes along the time course of 12 example bouts (colored according to their change in heading angle). Post-bout position and orientation are marked by black dots and arrows, respectively. The trajectory of the midpoint between the eyes forms a circular arc, with its length determined by the change in heading angle. These arcs matched the interpretation of rotation around *y*_*rot*_. The theoretical lines, representing the optional positions for each selected change in heading angle, are colored according to the change in heading angle. **C:** The trajectory of the midpoint between the eyes along the time course of bouts with a similar change in heading angle (40±3 degrees, 92 trajectories) (trajectories of different fish are indicated in different colors). Post-bout position and orientation are marked by black dots and arrows, respectively. Bouts of similar changes in heading angle show similar trajectories in the first phase of the movement (similar arcs). These bouts are completed with different distance traveled along the theoretical line (orange line) defined by the change in heading angle. **D:** The rotation point moves with the increase in body size. Two bigger black dots represent the two example fish shown in E. **E:** Example two fish (7 and 15 dpf fish represented by the two bigger dots in panel 3D), showing that the rotation point is positioned at the bottom edge of the swim bladder regardless of the size of the fish.

Additionally, the model suggested that after rotating larvae move forward or backward along the predicted linear line defined by the change in heading angle. We collected all bouts in the dataset with a particular change in heading angle (40°± 3°, n=92), and examined the trajectory of the point between the eyes along the time course of the movement as well as the outcome of these bouts. For this change in heading angle, 84% of post-bout positions were aligned along the predicted line with an error smaller than 0.2 mm, showing fish moving forward or backward after the rotation on the same predicted line (Figure 3C). This movement implementation was robust across other changes in heading angle (Figure S4). Together, movements are thus composed of two components: a rotation around a rotation point, and a movement forward or backward along a linear line defined by both the change in heading angle and the rotation point.

We then examined the position of the rotation point along the fish body axis. We assumed that the rotation point would be consistent in terms of its relative position for fish of different sizes. Therefore, we expected *y*_*rot*_ to increase with fish length. Since the variance in fish length in our dataset was rather low due to the narrow range of fish ages (5-7 dpf), we added to the dataset 13 older fish (14-15 dpf). The model fit the outcomes of these 14-15 dpf fish with similar performance to that for the 5-7 dpf fish (Figure S4). To identify the position of the rotation point on the body axis, we applied the model separately for each fish, extracting *y*_*rot*_ for each of our 31 fish (See Methods). The distance from the midpoint between the eyes to the rotation point (*y*_*rot*_) increased with fish length (Figure 3D). The rotation point was positioned at the bottom edge of the swim bladder across fish lengths (Figure 3E), suggesting that its relative position is preserved across individuals.

### The outcome repertoire explains the change in heading angle given a prey angle

Next, we used these insights into the repertoire of movement outcomes and their corresponding movement implementations to explain how fish interact with their prey. In particular, we sought to determine the guiding rule by which fish select the change in their heading angle for a given prey. To generate a robust dataset to study the interaction with the prey during the hunt, we collected all hunting events that were successfully completed, ensuring the identity of the target was unambiguous. In these events, we selected bouts for which we had access to the pre- and post-bout prey angles (1248 bouts in 300 events). The prey angle is usually measured relative to the point on the head which is the midpoint between the eyes 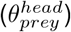 ^7,18,37^ and this prey angle is considered to elicit a particular change in fish heading angle (Δ*θ*_*head*_) (Figure 4A). Specifically, the relation between this prey angle and the change in fish heading angle was 0.58 ± 0.02 (Figure 4B), in agreement with previous reports ^7,37^. This suggests that fish do not fully close the angle to the prey with each movement, but reduce the angle to the prey by 60%. However, on examining the relation between this prey angle before and after each movement during the hunt, we found that fish maintain their prey angle within the ± 20° range throughout the hunting event (Figure 4C). This indicates that the fish orient themselves exactly towards the prey, rather than closing only 60% of the prey angle.

**Figure 4:**
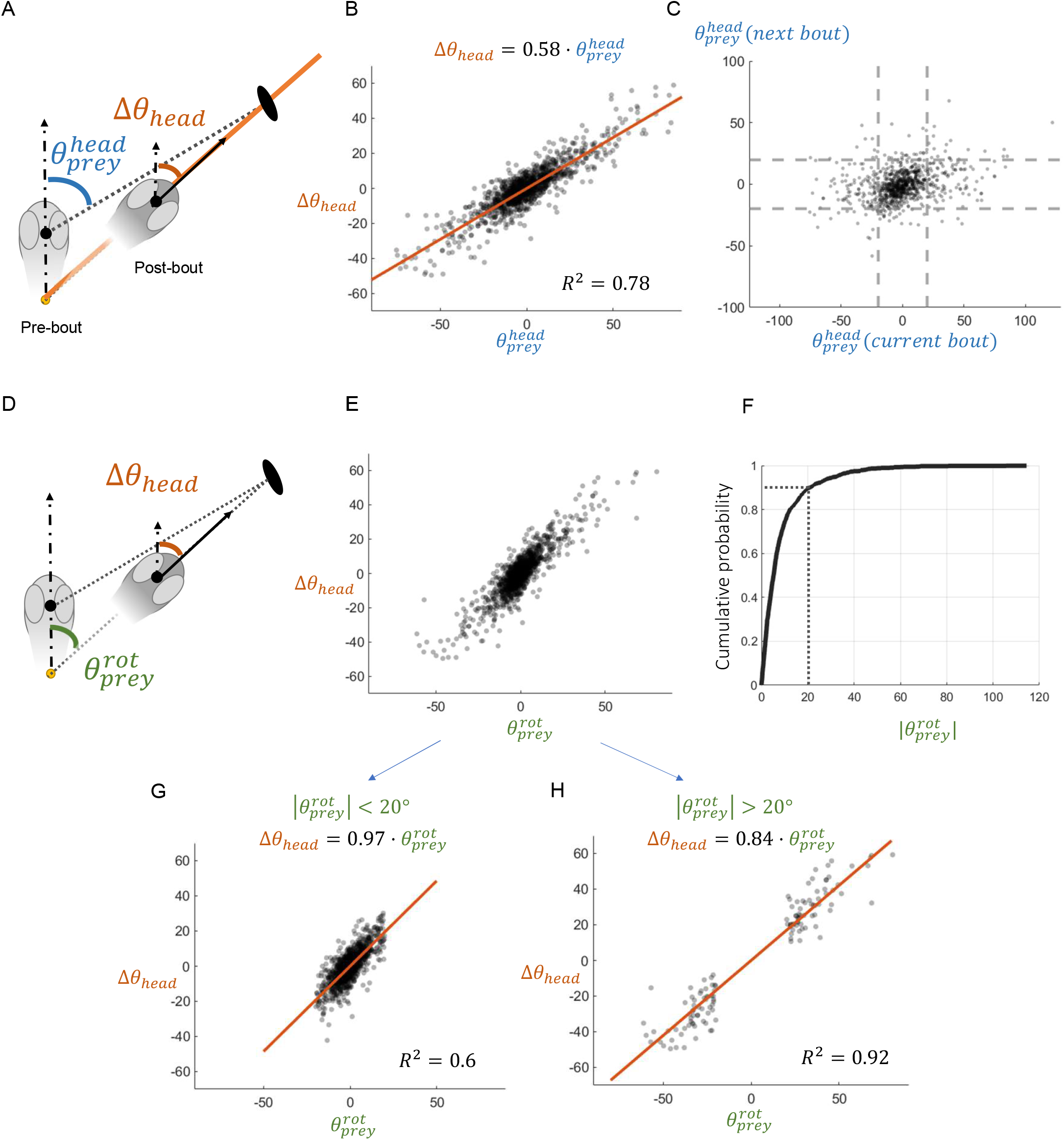
The change in heading angle matches the prey angle. **A:** A schematic of the interaction of the fish with the prey. Pre-bout head position is marked in bright grey head, with a dot representing fish position and a dashed arrow representing pre-bout heading direction. The prey angle 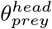 is measured relative to the pre-bout position. The reactive post-bout outcome is marked in dark grey fish head, with a black dot and an arrow for position and change in heading angle, respectively. The dashed arrow represents the pre-bout heading angle. Note that for another prey with the same prey angle (i.e. on a different position along the dashed line), the fish will still be positioned on the orange line. Therefore, if fish rotate in 60% of the prey angle, they may not face the prey following the movement. **B:** The relation between the change in heading angle and the prey angle 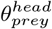 when measured relative to the midpoint between the eyes is characterized by a slope of 0.58. This suggests that the change in heading angle is 58% of the prey angle. **C:** Prey angles before and after movements show that in the majority of cases, the post-bout prey angle remained in the *±*20° range, as indicated by the dashed lines. The relation between the pre- and post-bout prey angle showed a slope of 0.17 (*R* = 0.3, *p value <* 1*e −* 20). **D:** A schematic of the interaction of the fish with the prey (similar to panel A). The prey angle is measured with respect to the rotation point 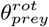 rather than the midpoint between the eyes. Note that the change in heading angle with respect to the midpoint between the eyes is identical to the change in heading angle when measured with respect to the rotation point. **E:** The relation between the change in heading angle and the prey angle when measured relative to the rotation point shows two regimes. **F:** Cumulative distribution of the prey angle when measured relative to the rotation point. 90% of prey angles are smaller than 20 degrees (in absolute value). **G:** For prey angles smaller than 20 degrees, the change in heading angle matches the prey angle. This suggests that fish fully orient themselves towards the prey. **H:** For prey angles larger than 20 degrees (comprising 10 % of prey angles), fish closes most (84%) of the angle to the prey.

To address this inconsistency we first computed the prey angle relative to the uncovered rotation point 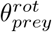 rather than the point on the head (Figure 4D). The change in heading angle remained identical irrespective of the reference point (mid point between the eyes or the rotation point) utilized for the measurement (Figure 4D). We then examined the relation between this prey angle and the selected change in the heading angle for each bout (Figure 4E). The overall relation was measured at 0.87 ± 0.03 (Figure S4A) but primarily clustered into two regimes. For 90% of prey angles, ranging from *−*20° to 20° (Figure 4F), the relation between prey angle and the change in heading angle was 0.97 ± 0.05 (Figure 4G). For 10% of prey angles, the relation of the change in heading angle was smaller and on average reached 0.84 ± 0.04 (Figure 4H). These results suggest that for most bouts during the hunting event, fish rotate around their axis to fully close the prey angle, aiming to place the prey in front of them.

Together, by examining the behavior with respect to the uncovered rotation point, we provide a substantially different account of the larval hunting behavior. If the fish close 60% of the prey angle but are free to select different positions along the linear line, there is a wide range of possible post-bout prey angles. Therefore, to place the target in front of them, larvae are limited to certain positions, thus coupling the selection of orientation and position. We thus suggest an alternative account: the fish first close the entire angle to their target, placing their prey in front of them. Following this rotation, fish move forward or backward while maintaining the same close-to-zero prey angle for all positions along the linear line. Therefore the selection of distance is decoupled from the change in heading angle (Figure S5F). On average, the observed 60% change in heading angle can be geometrically explained by the relation between the prey angle with respect to the midpoint between the eyes and the prey angle with respect to the rotation point (as shown on our data Figure S5E).

### The change in heading angle for a given prey is dependent on prey distance

The relation between the prey angle (with respect to the rotation point) and the change in heading angle revealed that in most cases fish close the entire angle to their target prey. However, for a small subset of cases, the change in heading angle was on average smaller than the entire prey angle. We thus sought to determine what prey features can potentially drive a smaller change in heading angle. More particularly, we asked whether a large distance to the prey or a large prey angle (as suggested by Figure 4H) affects this selection (Figure 5A). Fish interacted with targets at 1 mm-5 mm from their rotation point, with 71% of prey distances being less than 3 mm (Figure 5B). We examined the relation between the prey angle and the change in heading angle, labeling the change in heading angle according to the prey distance (Figure 5C). We observed smaller changes in heading angle for more distant targets (a slope lower than 1) and a complete closure of the angle to the prey (a slope of 1) for short-distanced prey (Figure 5C), suggesting that the relation between the change in heading angle and the prey angle, for a small fraction of movements, is distance-dependent.

**Figure 5:**
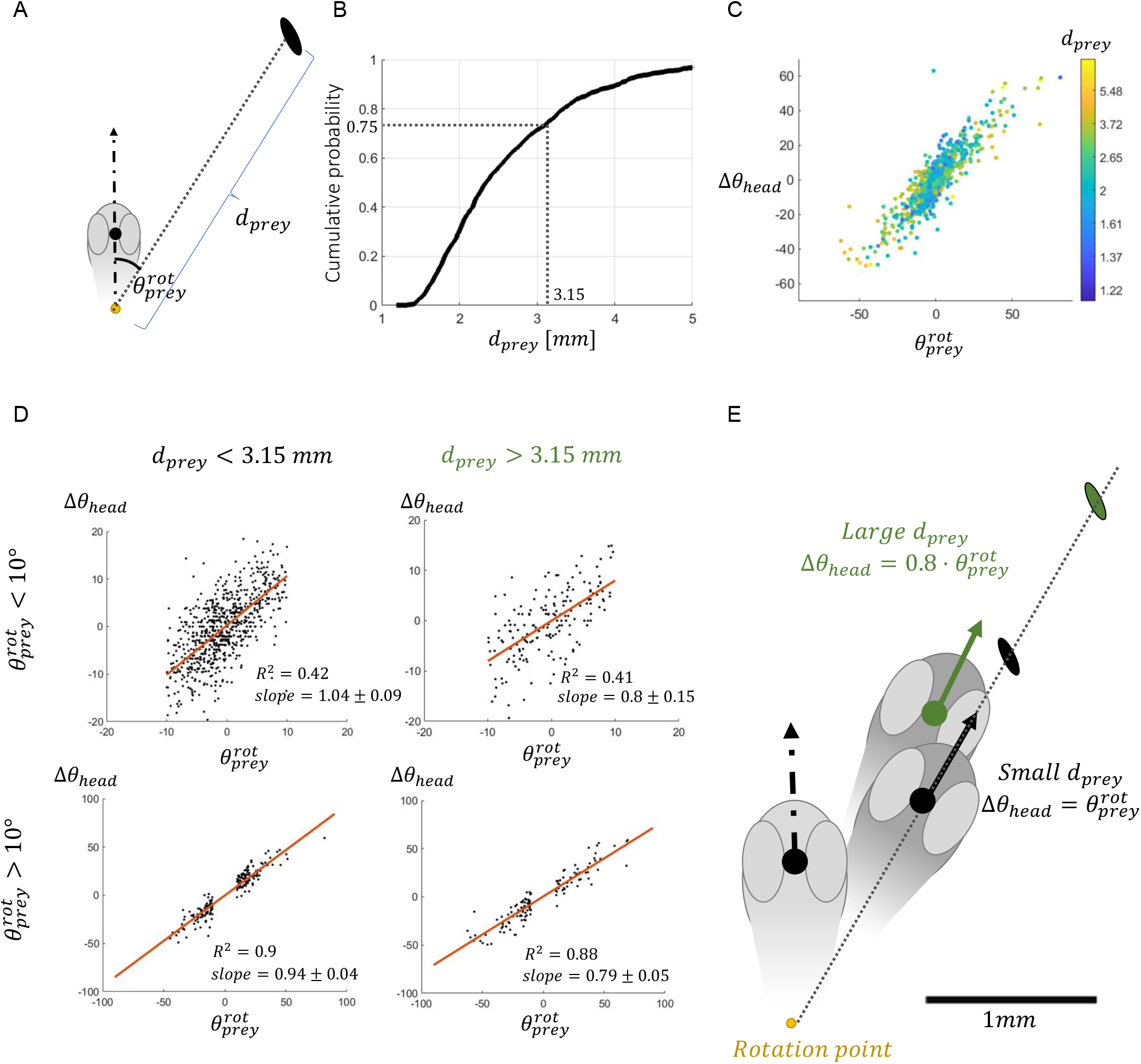
The change in heading angle in response to a prey item depends also on prey distance. **A:** A schematic of prey angle and distance, calculated relative to the rotation point. **B:** A cumulative distribution of prey distance during the hunting event. 75% of prey distances are below 3.15 mm. **C:** The relation between the change in heading angle and the prey angle measured with respect to the rotation point for each bout. Each bout is colored according to the prey distance. Short-range prey distances show a larger slope compared to long-range prey distances. **D:** Prey distance, and not prey angle, affects the slope between prey angle and the change in heading angle. The bouts were split according to the pre-bout small and large prey angle (upper and lower rows) and into pre-bout small and large prey distance (left and right columns) based on the 75th percentiles of both measures (10 degrees for prey angle and 3.15mm for prey distance, see Figure 4F and Figure 5B). For short-range prey distance the relation between prey angle and turn angle was 1. This was robustly similar for small and large prey angles, as indicated by overlapping confidence intervals of the slopes (left column). For long-range prey distances, this relation was reduced to 0.8, and remained robust across the different prey angles (right column). **E:** The suggested algorithm for the selection of change in heading angle given prey angle and distance. For short distances, fish rotate around their rotation point and align themselves directly towards the prey. Subsequently, fish move forward/backward along the line determined by the change in heading angle. For large prey distance, fish rotate around their rotation point and turn 80% of the prey angle.

To study the relation between prey angle, prey distance, and the selected change in heading angle, we split our data into short and long-range prey distances and small and large prey angles based on the 75th percentiles of both parameters (Figure 4F, Figure 5B). When prey were closer, fish changed their heading angle similarly for both small and large prey angles, aligning their heading angle to match the entire prey angle (Figure 5D, left column). However, for any given prey angle, they selected smaller changes in heading angle for the long-range distances compared to short distances (Figure 5D, left column versus right column). These results were robust also when dividing the data according to the 50th or 90th percentiles (Figure S6). Together, these findings show that for short-range prey distances, fish rotate around their rotation point, close the prey angle, and orient themselves in front of the prey. However, for long-range prey distances, fish rotate only partially and close most but not all of the angle to the target (Figure 5E). These results suggest that the change in heading angle depends not only on prey angle but also on prey distance.

## Discussion

Much like in a board game, it is essential to master the repertoire of movements each piece can make, as this forms the foundation for understanding the overall strategy. The same rule applies when studying the larval zebrafish hunting behavioral algorithm. We studied the possible changes in fish position and orientation following a single movement, which we termed movement outcomes. We show that the repertoire of outcomes can be fully described using three parameters, two positional parameters and an orientation parameter. We mathematically uncovered the relationship between position and orientation and thereby reconstructed the complete repertoire. This relation suggested that the position of larvae after a movement accurately predicts the change in heading angle. In addition, a particular change in heading angle establishes a limited set of positions aligned on a linear line in space. This uncovered repertoire unraveled a fundamental rule guiding the interaction between the larval zebrafish and its target. We show that given a prey, in most movements larvae will turn the entire prey angle, placing themselves on a particular linear line in front of the prey. In addition, for distant paramecia, both the prey angle and distance dictate the change in heading angle. These results provide a complete and continuous description of the repertoire of movements and its dimensionality. The uncovered nature of the repertoire uncovered the underlying algorithmic rules of the behavior and sheds light on its neural implementation.

Based on the relation between position and orientation, our work suggested a two-dimensional outcome repertoire. Therefore, the repertoire of all available positions and orientations following a single movement is fully described using two parameters. We showed that the specific pair of azimuthal angle and distance can capture the outcome unambiguously (Figure 2B-D). Other pairs of parameters (such as change in heading angle and azimuthal angle, or change in heading angle and bout distance) were insufficient to capture certain movements in the repertoire (Figure S3). An alternative option for quantifying the outcome is based on the movements principles uncovered by the repertoire, and suggests using the change in heading angle and the distance between the pre- and post-bout rotation points (Δ*θ*_*head*_ and *d*^*′*^) as the parameters that describe the repertoire (Figure 3A). This alternative may appear similar to the change in heading angle and distance measured from the point on the head (midpoint between the eyes) (Δ*θ*_*head*_ and *d*), which we found to be insufficient. However, this alternative is distinct because *d*^*′*^ is the distance traveled along the linear line defined by the change in heading angle, ensuring a single position and orientation for each pair of values. These two alternative approaches to quantifying an outcome (*ϕ*_*az*_ and *d* relative to the point between the eyes or Δ*θ*_*head*_ and *d*^*′*^ relative to the rotation point) are equivalent.

The low dimensionality of the repertoire of outcomes observed in this study is not surprising. To begin with, in experiments conducted in shallow water, outcomes are bounded by at most three dimensions. Nevertheless, the fact that the repertoire of outcomes is two-dimensional has several important implications. First, as we showed here, moving from three to two dimensions implies a structure to the repertoire. This structure uncovered a fundamental movement principle and a guiding rule for action selection. Second, the low dimensionality of the outcome repertoire has implications for our understanding of the movement repertoire. More particularly, the high-dimensional movements and the low-dimensional outcomes offer two alternatives for interpreting the dimensionality of movements. The first option suggests a potential low-dimensional structure of the movements as well. In this case, every movement leads to a unique outcome, and choosing the outcome is equivalent to choosing the movement. Alternatively, if movements are high-dimensional then different movements lead to a similar outcome. This suggests a more complicated action selection mechanism: animals either select a high-dimensional movement (from which the outcome is derived), or implement a two-stage mechanism, selecting a desired outcome and subsequently selecting a specific movement to implement it. While the relation between outcomes and movements is still unknown, one way to bridge this gap is by using the trajectories of the midpoint between the eyes shown in Figure 3C. These trajectories can be thought of as outcomes of a single movement over time, and our results suggest a potential link between the final outcome, outcome over time, and movements. Overall, while the dimensionality of tail movements remains a fundamental open question, the notion of outcomes can aid at uncovering the dimensionality of movements. Therefore, further work focused on the relation between tail movements and outcomes is required.

The low-dimensional nature of the outcome repertoire has further implications for neural information processing. Visual information coming from the outside world during the hunt is high-dimensional, and includes the target prey, its size, angle, velocity, and other prey items in the visual field of the fish. If actions are selected based on their outcome, this complex detailed visual information should eventually be processed to select the best outcome for a given goal. High-dimensional sensory information and a low-dimensional outcome repertoire suggest that the visuomotor transformation may compress the world into low-dimensional actions. This further suggests that the visuomotor transformation process may be invariant to several features of the external world. Supporting evidence for such a transformation comes from recordings of hindbrain neural activity in fish moving their tail and eyes, showing that most behavior-related population activity can be captured using two dimensions ^14^. This experiment, like our work, was conducted in a two-dimensional environment, and their reported two-dimensional neural activity corresponding to the behavior aligns with our two-dimensional outcomes. While this finding raises questions regarding the dimensionality of movements, it aligns with the idea of two-dimensional outcomes.

An interesting open question is whether fish select a desired outcome before implementing the movement, or respond to the visual field as they move and therefore modify their movement while moving (the latter option suggests that fish can parse the visual scene while moving). On the one hand, our work suggests that fish first turn to face their prey and then move forward or backward (Figure 3A). This allows the fish to engage in an online decision making: keep rotating until they see the prey, and move forward or backward relative to the observed prey. On the other hand, recent work has suggested that visual processing is suppressed during self-motion by motor-related inhibition ^1^. In addition, the trajectories of the midpoint between the eyes for the different changes in heading angle (Figure 3B) show that while the fish turn and then move forward or backward, in some movements they start to progress forward or backward before they complete the entire turn angle (Figure 3B-C), while still ending up on the line suggested by the model of positions given a change in heading angle. This implies that before the rotation phase is completed, the fish aim toward the theoretical line of possible positions for this particular change of heading angle. While the low-dimensional repertoire of outcomes calls for an in-advance action selection process, further evidence supporting online or in-advance processes is required from both a behavioral and a neuronal point of view.

Our work empirically uncovered a rotation point based on the relation between the change in heading angle to the positional parameters (equation in Figure 2C, and Figure 3, see also Methods). Using this point we were able to better explain both the structure of the repertoire and the behavior. We showed that the rotation point remains at the bottom edge of the swim bladder regardless of the size of the fish. Fish are expected to rotate around their center of mass, which depends on the distribution of mass along the fish body. If indeed the rotation point is the center of mass, we can make further predictions as to the repertoire of outcomes in three-dimensional environments known to be utilised in naturalistic behaviors ^20^. In the general case, the outcome of a movement in a three-dimensional environment should be represented by 5 variables (2 for fish orientation, 2 for direction of movement, 1 for distance). If fish rotate around a single point, then outcome can be simplified into a three-dimensional representation (2 variables to represent the rotation around the rotation point, and 1 for the distance travelled following the rotation).

Previous works have suggested that fish turn about 60% of the prey angle ^7,37^ and further indicated that fish cut off 50% of the prey angle with each bout ^7^. These observations may suggest that the fish may not be facing the target during parts of the hunt. However, our work suggests that fish turn the entirety of the prey angle, positioning the target directly in front of them. The previously reported partial rotation toward the prey and the fact fish are facing the prey after every movement are not contradictory; they instead arise from different reference points used to measure the prey angle. The prey angle can be measured either with respect to the point between the eyes or the rotation point (Figure 5A,D). Since fish observe the world with their eyes and rotate around a separate rotation point, a correction is necessary when translating the observed prey angle into a selected change in heading angle. There are two possible ways to implement this correction. The first alternative is to use the observed prey angle and distance to calculate the prey angle relative to the rotation point, though this requires complex geometric computation. The second alternative relies on the observation that, on average, prey angles relative to the rotation point are smaller by 60% compared to the angles relative to the head (Figure S5E), using this as a constant factor. While this is an approximation, it results in relatively small errors in the desired rotation angle. Overall, while distinct interpretations to a turn of 60% of the prey angle were suggested, our model identified a reference point that revealed that the algorithmic rule of the fish is to turn the entirety of the prey angle.

While uncovering the repertoire has allowed us to identify a fundamental rule underlying the behavioral algorithm of hunt, what determines the selection of distance and its relation to prey parameters remains an open question. Our model suggests that the choice of bout distance relative to the rotation point is independent of selecting the change in heading angle, since all possible positions ensure that the prey angle is near zero. This was further supported by the observation that, for a given change in heading angle, all possible distances are observed (Figure S5F). With the comprehensive capturing of the repertoire, we are now better equipped to characterize the factors that shape distance selection. Particularly, when measuring bout distance relative to the midpoint between the eyes, some of the measured distance is the result of the rotation itself (as illustrated by the blue line in Figure S5F). The relation between bout distance given a prey distance provided, on average, a linear relation between prey distance and bout distance ^7^ (Figure S6C). However, our data show that despite this generally linear relation, the selection of bout distance given the prey distance is highly variable (Figure S6D). Further work is thus required to uncover the guiding rule underlying the selection of bout distance during the hunt.

The relation between the change in heading angle and the prey angle with respect to the uncovered rotation point suggests that for most movements, the fish turn the entirety of the prey angle to place the target in front of them. We further showed that this change in heading angle depends not only on prey angle but also on prey distance, with large prey distance eliciting a smaller turn. The relation between prey angle, prey distance, and turn angle is shown in Figure 5C. Long-range distance prey (yellow dots) exhibited a smaller slope compared to the short-range distance prey (blue points). In our analysis, we divided the dataset into two subsets: with 90% and 10% of the bouts respectively obeying relationships of 0.97 and 0.84. While we suggest a slope of 1 for 90% of the data and 0.84 for the remaining 10%, there may be a more gradual change in slope as a function of prey distance (Figure 5C). Further work is required to fully understand the effect of prey distance on the selected change in heading angle.

## Methods

### Zebrafish maintenance

Nacre zebrafish (Danio rerio) embryos expressing elavl3:H2B-GCaMP6s^38^ were collected and raised according to established procedures ^23^ and raised at 27°C with a light and dark cycle of 14/10 h. Larvae were fed live rotifers (Brachionus plicatilis) daily from 5 dpf. All procedures were performed with approval from The Hebrew University of Jerusalem Animal Ethics Committee.

### Imaging natural hunting behavior of freely swimming larvae

Single larvae, 5-7 dpf (n=18), were placed in a transparent bottom plate (20 mm diameter and 2.5 mm deep) with 30 Paramecia for 15 minutes (CoverWell Imaging Chambers, Catalogue number 635031, Grace Biolabs). The plate was illuminated from above with white visible light and Infra-red LEDs were placed beneath the plate (L850 nm, LDR2-100IR2-850-LA powered by PD3-3024-3-PI, CCS Inc., Kyoto, Japan). Room temperature was maintained at 27°C. Natural larvae behavior was captured from above, using a high-speed camera (MIkrotoron 4CXP), imaging at 500 fps and 45 microns per pixel.

To study the position of the rotation point as a function of fish length (Figure 3D), we recorded 13 fish at the ages of 14-15 dpf under the same protocols.

### Tracking fish and prey features

The recordings of the natural hunting behavior were analyzed post-hoc using an algorithm we developed in the lab based on computer vision and graph theory techniques. This custom-made algorithm detected the fish, extracted its features, identified time points of movements, and detected prey items and their trajectories.

### Fish detection

To detect the fish and extract its features, we isolated the fish from its environment on a frameto-frame basis, thus eliminating any interference from prey or noise (dust particles in the dish). We removed the contour of the imaging dish from the frame by searching for contours with a convex hull forming a large elliptic shape. The removal of the dish contour made the fish the largest object in the frame, which we verified based on its the overall shape and size.

### Fish feature extraction

After identifying the fish object, we computed the contour around it and searched for objects that reside within the specified region marked as the fish. We defined the eyes as the two darkest objects within the fish object, which are elliptical, adjacent, and falling within a certain size range. Fitting an ellipse to each of the eye contours provided both the ellipse center and orientation as expressed by its major and minor axes. We then constructed the head direction vector by setting the origin point of this vector as the midpoint between the eye centers, and the head direction as a vector orthogonal to the axis connecting the two eye centers. The relative angle between the fish heading and the major axis of the eye-fitted ellipse was used to evaluate eye convergence and divergence. In the young dataset (5-7 dpf), 939 events were identified, including 262, 177, 500 hit, miss, and abort events respectively, and overall 4006 bouts. In the older dataset of 14-15 dpf 636 events were detected including 274, 146, 216 hit, miss and abort events, and overall 2545 bouts.

### Automatic identification of hunting events

We used eye convergence and divergence as the markers for the onset and offset of the hunting events ^5,6,29,34^. Based on the statistics of eye angles in a training set of our data, we identified an eye angle threshold above which the frame was classified as a frame during the hunt. This threshold was used to detect the onset and offset of each event in the data. A sequence of frames of detected converged eyes was suggested as a potential event if it occupied at least 120 consecutive frames while combining segments which were separated by less than 10 frames. This automatic machinery generated a list of potential hunting events, marked by their onset and offset frame, and these suggested events were manually inspected and validated.

### Bout detection

To identify the frames in which the fish performed bouts of movement, we used our tracking of the fish contour and the fish midline. We selected a set of equidistant points along the tail midline, ordered from the tail tip to the fish head. We then calculated the tail tip velocity as the change in Euclidean distance between consecutive frames. We set a threshold for tail velocity above which the fish was performing a movement, defining the minimal duration of a bout as 5 frames.

### Extracting bout outcomes and the relation to prey

#### Quantifying bout outcomes

Outcomes were quantified using three parameters. Bout distance (*d*), bout change in heading angle (Δ*θ*_*head*_), and bout azimuthal angle (*ϕ*_*az*_) (Figure 1B,C). Larval position was defined as the middle point between the detected eyes (Figure 1A,B). Bout distance was set as the Euclidean distance between the larval pre- and post-bout position (Figure 1B,C). The change in azimuthal angle was defined as the angle between the pre-bout heading direction and the fish movement direction. Movement direction was defined by larval pre- and post-bout position (Figure 1B,C). The change in heading angle was defined as the difference between pre- and post-heading direction (Figure 1B,C).

#### Quantifying prey features

Prey features that were considered were prey distance and prey angle. Prey distance was defined as the Euclidean distance between the prey center of mass and the fish position. Prey angle was defined as the angle between the heading direction and the direction defined by fish position and prey position (Figure 4A).

#### Evaluating of the rotation point

To evaluate the position of the rotation point *y*_*rot*_, we fitted the model to all bout outcomes based on the equation shown in Figure 2C. The model uncovered the relation between the three outcome parameters (*d*, Δ*θ*_*head*_, *ϕ*_*az*_), and was based on a single parameter *y*_*rot*_. The fit identified *y*_*rot*_ that minimized 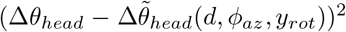, where 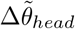 is the predicted change in heading angle based on the position parameters and on *y*_*rot*_. Estimating *y*_*rot*_ for all fish, we minimized the above prediction error quantity for all bouts together. When evaluating *y*_*rot*_ per fish (Figure 3D), we performed this minimization for the bouts of each fish separately. The per-fish variance of *y*_*rot*_ estimation (Figure 3D) was calculated by splitting all bouts of a single fish into 5 groups with equal number of bouts and fitting *y*_*rot*_ separately for each group. This provided 5 estimates of *y*_*rot*_ per fish and these estimates were used to calculate the variance.

#### Reconstruction of the empiric repertoire

The reconstruction of the repertoire in Figure 2D and S2 was performed based on the fits of all outcomes in the dataset using the model shown in Figure 2C. The fit of all outcomes in the 5-7 dpf dataset suggested that the rotation point is positioned 1 mm behind the midpoint between the eyes. The reconstruction was performed in two equivalent ways. For the first way, we used the equation in Figure 2C and calculated the change in heading angle given the distance and the azimuthal angle. For the second way, based on the interpretation presented in Figure 3A, we calculated the distance and azimuthal angle based on the change in heading angle and the distance relative to the rotation point (Δ*θ*_*head*_, *d*^*′*^). For each change in heading angle, we used distances ranging from a pre-defined initial value to 2.5 mm. This initial value for each change in heading angle was determined by fitting a second degree polynomial to the observed relation between Δ*θ*_*head*_ and *d*^*′*^

## Acknowledgements

We thank Haim Sompolinsky for valuable discussions. We thank Israel Nelken, Yoram Burak, and Mati Jushua for their comments on earlier versions of the article. We thank the Avitan lab members for their valuable feedback and discussions. We gratefully acknowledge funding from Israel Science Foundation grants 1684/20.

## References

1. Ali, M. A., Lischka, K., Preuss, S. J., Trivedi, C. A., and Bollmann, J. H. (2023). A synaptic corollary discharge signal suppresses midbrain visual processing during saccade-like locomotion. Nature Communications, 14, 7592.

2. Anderson, D. and Perona, P. (2014). Toward a science of computational ethology. Neuron, 84, 18–31.

3. Avitan, L., Pujic, Z., Mölter, J., McCullough, M., Zhu, S., Sun, B., Myhre, A.-E., and Goodhill, G. J. (2020). Behavioral signatures of a developing neural code. Current Biology, 30, 3352–3363.e5.

4. Berman, G. J., Choi, D. M., Bialek, W., and Shaevitz, J. W. (2014). Mapping the stereotyped behaviour of freely moving fruit flies. J. R. Soc. Interface, 11, 20140672.

5. Bianco, I., Kampff, A., and Engert, F. (2011). Prey capture behavior evoked by simple visual stimuli in larval zebrafish. Frontiers in Systems Neuroscience, 5.

6. Bianco, I. H. and Engert, F. (2015). Visuomotor transformations underlying hunting behavior in zebrafish. Current Biology, 25, 831–846.

7. Bolton, A. D., Haesemeyer, M., Jordi, J., Schaechtle, U., Saad, F. A., Mansinghka, V. K., Tenenbaum, J. B., and Engert, F. (2019). Elements of a stochastic 3d prediction engine in larval zebrafish prey capture. eLife, 8, e51975.

8. Borla, M. A., Palecek, B., Budick, S., and O’Malley, D. M. (2002). Prey Capture by Larval Zebrafish: Evidence for Fine Axial Motor Control. Brain Behavior and Evolution, 60, 207–229.

9. Budick, S. and O’Malley, D. (2000). Locomotor repertoire of the larval zebrafish: swimming, turning and prey capture. Journal of Experimental Biology, 203, 2565–2579.

10. Burgess, H. A. and Granato, M. (2007). Modulation of locomotor activity in larval zebrafish during light adaptation. Journal of Experimental Biology, 210, 2526–2539.

11. Costa, A. C., Ahamed, T., Jordan, D., and Stephens, G. J. (2024). A markovian dynamics for caenorhabditis elegans behavior across scales. Proc. Natl. Acad. Sci. U. S. A., 121, e2318805121.

12. Egnor, S. R. and Branson, K. (2016). Computational analysis of behavior. Annual Review of Neuroscience, 39, 217–236.

13. Ehrlich, D. E. and Schoppik, D. (2019). A primal role for the vestibular sense in the development of coordinated locomotion. eLife, 8, e45839. eLife 2019;8:e45839.

14. Feierstein, C. E., de Goeij, M. H., Ostrovsky, A. D., Laborde, A., Portugues, R., Orger, M. B., and Machens, C. K. (2023). Dimensionality reduction reveals separate translation and rotation populations in the zebrafish hindbrain. Current Biology, 33, 3911–3925.e6.

15. Filosa, A., Barker, A., Dal Maschio, M., and Baier, H. (2016). Feeding state modulates behavioral choice and processing of prey stimuli in the zebrafish tectum. Neuron, 90, 596–608.

16. Gahtan, E., Tanger, P., and Baier, H. (2005). Visual prey capture in larval zebrafish is controlled by identified reticulospinal neurons downstream of the tectum. Journal of Neuroscience, 25, 9294–9303.

17. Girdhar, K., Gruebele, M., and Chemla, Y. R. (2015). The behavioral space of zebrafish locomotion and its neural network analog. PLOS ONE, 10, 1–18.

18. Henriques, P. M., Rahman, N., Jackson, S. E., and Bianco, I. H. (2019). Nucleus isthmi is required to sustain target pursuit during visually guided prey-catching. Current Biology, 29, 1771–1786.e5.

19. Hernández, L. P., Barresi, M. J. F., and Devoto, S. H. (2002). Functional morphology and developmental biology of zebrafish: Reciprocal illumination from an unlikely couple1. Integrative and Comparative Biology, 42, 222–231.

20. Horstick, E. J., Bayleyen, Y., Sinclair, J. L., and Burgess, H. A. (2017). Search strategy is regulated by somatostatin signaling and deep brain photoreceptors in zebrafish. BMC Biology, 15, 4.

21. Johnson, R. E., Linderman, S., Panier, T., Wee, C. L., Song, E., Herrera, K. J., Miller, A., and Engert, F. (2020). Probabilistic models of larval zebrafish behavior reveal structure on many scales. Current Biology, 30, 70–82.e4.

22. Jouary, A. and Sumbre, G. (2016). Automatic classification of behavior in zebrafish larvae. bioRxiv, p. 052324.

23. M, W. (2007). The zebrafish book; a guide for the laboratory use of zebrafish (danio rerio).

24. Marques, J. C., Lackner, S., Félix, R., and Orger, M. B. (2018). Structure of the zebrafish locomotor repertoire revealed with unsupervised behavioral clustering. Current Biology, 28, 181–195.e5.

25. McClenahan, P., Troup, M., and Scott, E. K. (2012). Fin-tail coordination during escape and predatory behavior in larval zebrafish. PLOS ONE, 7, e32295.

26. McElligott, M. B. and O’Malley, D. M. (2005). Prey Tracking by Larval Zebrafish: Axial Kinematics and Visual Control. Brain Behavior and Evolution, 66, 177–196.

27. Mearns, D. S., Donovan, J. C., Fernandes, A. M., Semmelhack, J. L., and Baier, H. (2020). Deconstructing hunting behavior reveals a tightly coupled stimulus-response loop. Current Biology, 30, 54–69.e9.

28. Mirat, O., Sternberg, J., Severi, K., and Wyart, C. (2013). Zebrazoom: an automated program for high-throughput behavioral analysis and categorization. Frontiers in Neural Circuits, 7.

29. Muto, A. and Kawakami, K. (2013). Prey capture in zebrafish larvae serves as a model to study cognitive functions. Frontiers in Neural Circuits, 7.

30. Muller, U. K. and van Leeuwen, J. L. (2004). Swimming of larval zebrafish: ontogeny of body waves and implications for locomotory development. Journal of Experimental Biology, 207, 853–868.

31. Nath, T., Mathis, A., Chen, A. C., Patel, A., Bethge, M., and Mathis, M. W. (2019). Using deeplabcut for 3d markerless pose estimation across species and behaviors. Nature Protocols, 14, 2152–2176.

32. Patterson, B. W., Abraham, A. O., MacIver, M. A., and McLean, D. L. (2013). Visually guided gradation of prey capture movements in larval zebrafish. Journal of Experimental Biology, 216, 3071–3083.

33. Petrucco, L., Lavian, H., Wu, Y. K., Svara, F., Štih, V., and Portugues, R. (2023). Neural dynamics and architecture of the heading direction circuit in zebrafish. Nature Neuroscience, 26, 765–773.

34. Semmelhack, J. L., Donovan, J. C., Thiele, T. R., Kuehn, E., Laurell, E., and Baier, H. (2014). A dedicated visual pathway for prey detection in larval zebrafish. eLife, 3, e04878. eLife 2014;3:e04878.

35. Stephens, G. J., Johnson-Kerner, B., Bialek, W., and Ryu, W. S. (2008). Dimensionality and dynamics in the behavior of c. elegans. PLOS Computational Biology, 4, e1000028.

36. Thorsen, D. H. and Hale, M. E. (2005). Development of zebrafish (danio rerio) pectoral fin musculature. Journal of Morphology, 266, 241–255.

37. Trivedi, C. and Bollmann, J. (2013). Visually driven chaining of elementary swim patterns into a goal-directed motor sequence: a virtual reality study of zebrafish prey capture. Frontiers in Neural Circuits, 7.

38. Vladimirov, N., Mu, Y., Kawashima, T., Bennett, D. V., Yang, C.-T., Looger, L. L., Keller, P. J., Freeman, J., and Ahrens, M. B. (2014). Light-sheet functional imaging in fictively behaving zebrafish. Nature Methods, 11, 883–884.

39. Westphal, R. E. and OMalley, D. M. (2013). Fusion of locomotor maneuvers, and improving sensory capabilities, give rise to the flexible homing strikes of juvenile zebrafish. Frontiers in Neural Circuits, 7.

40. Wiltschko, A. B., Johnson, M. J., Iurilli, G., Peterson, R. E., Katon, J. M., Pashkovski, S. L., Abraira, V. E., Adams, R. P., and Datta, S. R. (2015). Mapping sub-second structure in mouse behavior. Neuron, 88, 1121–1135.

