## Supplementary material for "A complete account of the behavioral repertoire uncovers principles of larval zebrafish hunting behavior": Supp Info

Yoav Rubinstein[1], Maayan Moshkovitz[1], Itay Ottenheimer[1], Sapir Shapira[1], Stas Tiomkin[2], Lilach Avitan[1]
[1]Edmond and Lily Safra Center for Brain Sciences, The Hebrew University of Jerusalem, 9190401, Israel.
[2]Computer Science Department, Whitacre College of Engineering, Texas Tech University, TX, 79409, USA.

Supplementary information

A

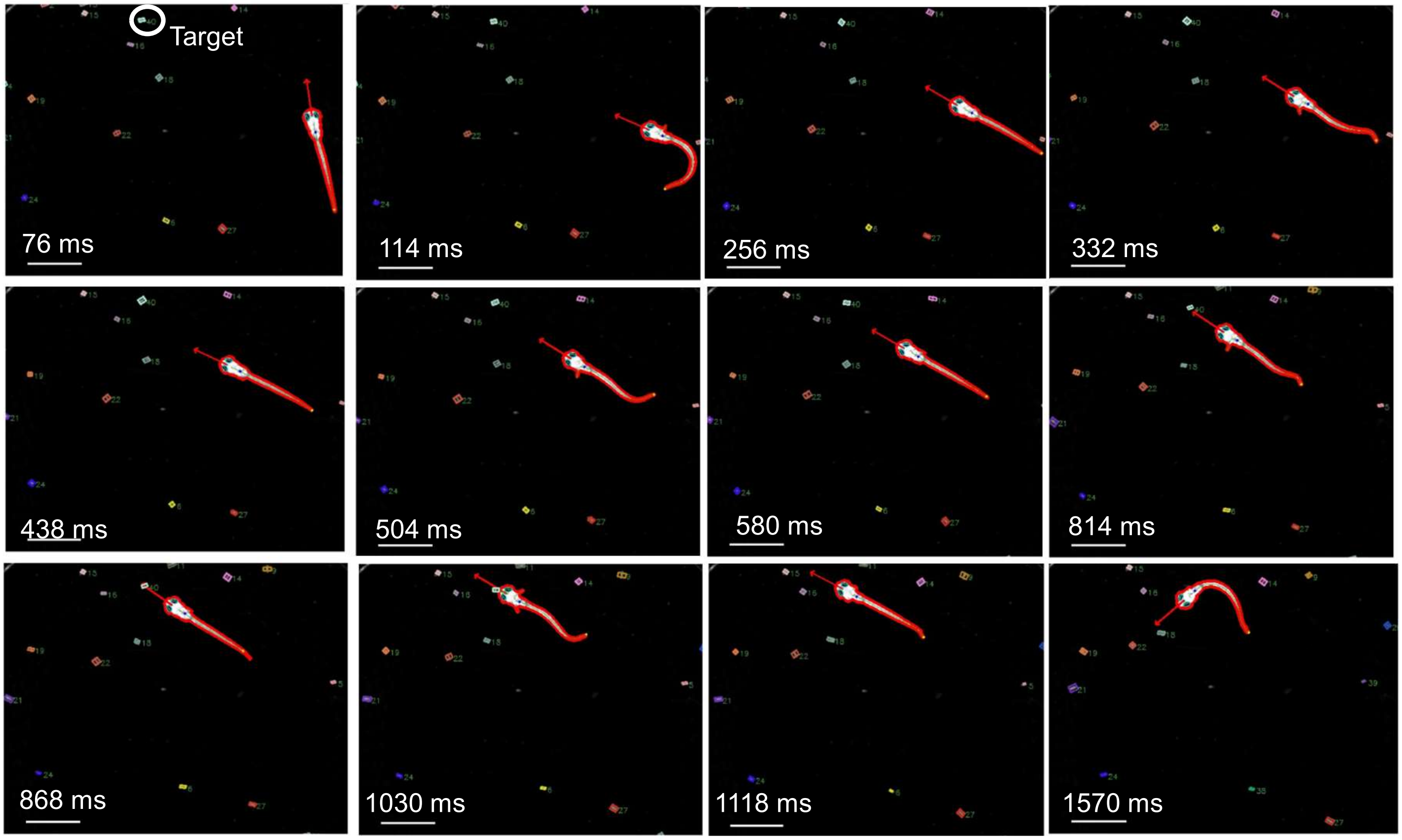


**Supplementary Figure 1. A larval zebrafish hunting event. A:** 12 example frames from a hunting event, tracked and annotated by our algorithm. The algorithm identified fish contour (red contour), heading angle (red arrow), swim bladder (blue point), tail midline (green line), tail tip (yellow point) and eyes (cyan). Paramecia (prey items) are marked by colored rectangles, where each paramecium is indexed. In this event, the larva uses 5 bouts to capture the prey, followed by an escape bout at the end of the event.

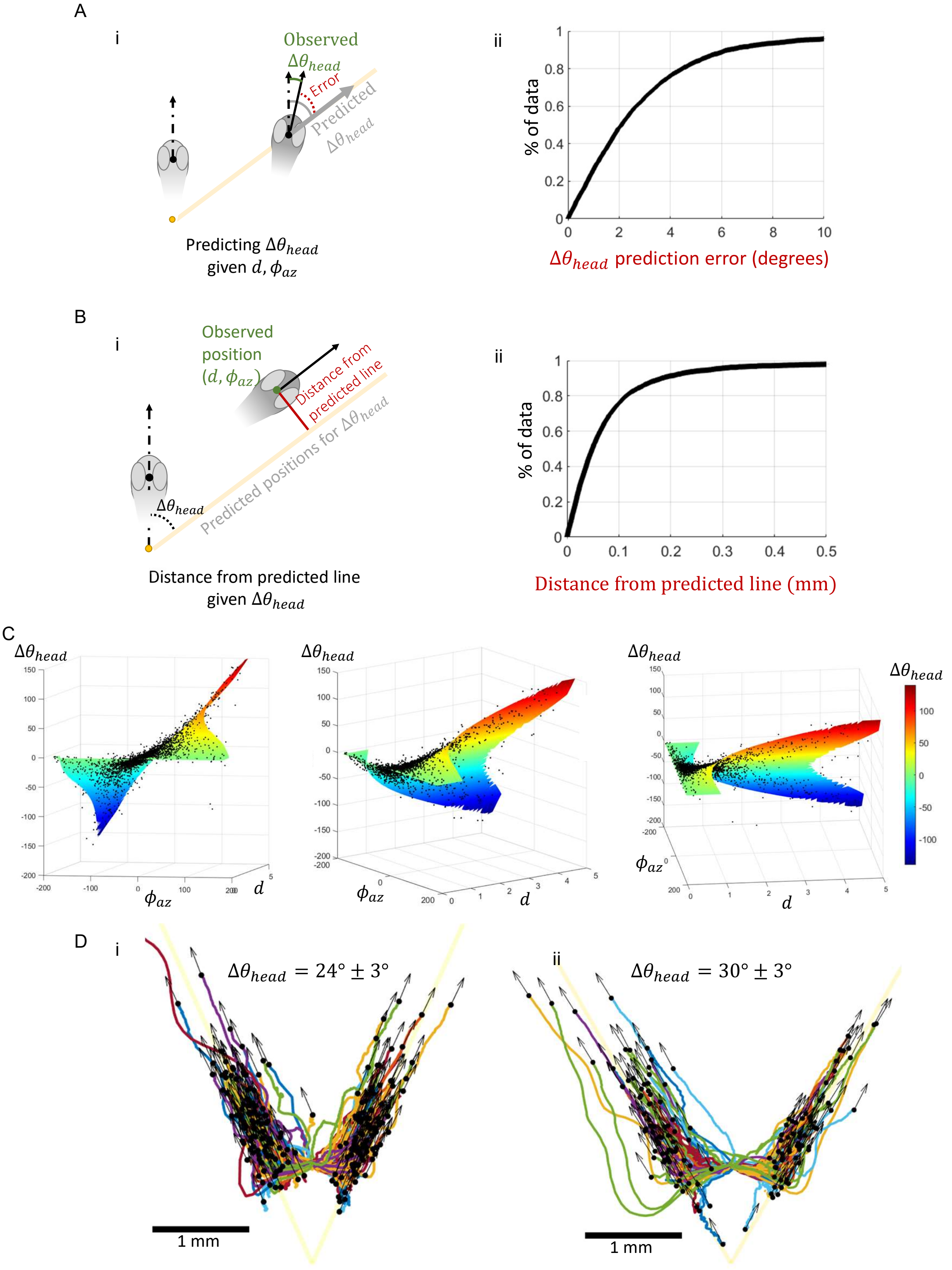

A
i
Observed
$\Delta\theta_{head}$
Error
Predicted
$\Delta\theta_{head}$
Predicting $\Delta\theta_{head}$
given $d, \phi_{az}$
ii
% of data
$\Delta\theta_{head}$ prediction error (degrees)
B
i
Observed
position
$(d, \phi_{az})$
Distance from
predicted line
Predicted positions for $\Delta\theta_{head}$
$\Delta\theta_{head}$
Distance from predicted line
given $\Delta\theta_{head}$
ii
% of data
Distance from predicted line (mm)
C
$\Delta\theta_{head}$
$\phi_{az}$
$d$
D
i
$\Delta\theta_{head} = 24° \pm 3°$
1 mm
ii
$\Delta\theta_{head} = 30° \pm 3°$
1 mm

**Supplementary Figure 2. The model accurately captures the repertoire of outcomes. A:** (i) The pre-bout position is shown by a light grey fish head, with the head position and head direction indicated by the point on its head and a dashed arrow respectively. The post-bout position is shown by a dark grey fish head, with position and heading angle indicated by the point on the head and a solid arrow respectively. The dashed arrow on the dark grey head indicates the pre-bout heading angle. Given the post-bout position (a dot on the dark grey fish head), the predicted change in heading angle is marked in grey. The observed change in heading angle is marked in green and the prediction error in red. (ii) Given the position of the fish ($\phi_{az}, d$), our model predicted the change in heading angle with an accuracy of 4 degrees for 76% of the bouts. **B:** (i) pre- and post-bout head position and direction similar to panel A(i). The predicted error in fish position (red line) is calculated by measuring the distance between the actual post-bout position (green point) for a specific change in heading angle and the theoretical line (orange line) predicted by the model for that same change in heading angle. (ii) Given the change in heading angle, our model predicted the position with an accuracy of 0.2 mm for 91% of the bouts. **C:** The measured repertoire of outcomes (black dots) and the reconstructed repertoire by our model (surface colored by the heading angle). The repertoire of outcomes forms a two-dimensinal manifold embedded in the three-dimensional space of the outcomes parameters (change in heading angle, distance and azimuthal angle). The manifold is a screw-like surface, with non-trivial relations between the different parameters (detailed in Figure 2C). For the 3d shape of the manifold see Supplementary Movie 5. **D:** Outcomes and their respective trajectories of the point between the eyes for bouts of $24 \pm 3$ (i) and $30 \pm 3$ (ii) change in heading angle. Colors represent trajectories from different fish. The trajectories, final positions, and orientations of the bouts (black arrows) align with the theoretical line predicted by our model. For each change in heading angle (yellow lines in i and ii, representing turns of $30^\circ$ and $24^\circ$, 88% and 93% of post-bout positions were aligned along the respective predicted lines with an error smaller than 0.2 mm. This suggests that the larvae progress along the theoretical line and end up on it.

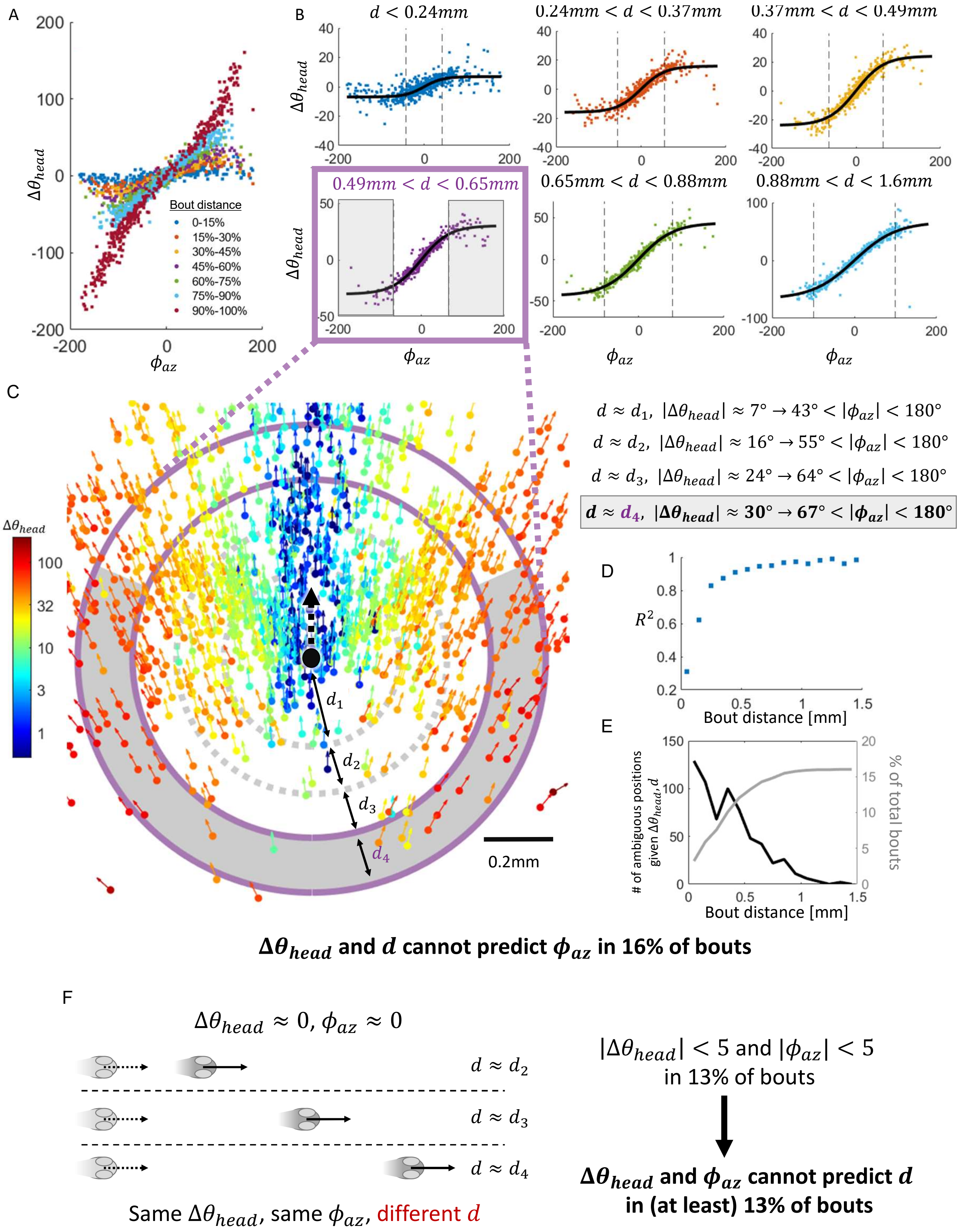

A
B
$\Delta\theta_{head}$
$\phi_{az}$
Bout distance
0-15%
15%-30%
30%-45%
45%-60%
60%-75%
75%-90%
90%-100%
$d < 0.24mm$
$0.24mm < d < 0.37mm$
$0.37mm < d < 0.49mm$
$0.49mm < d < 0.65mm$
$0.65mm < d < 0.88mm$
$0.88mm < d < 1.6mm$
C
$d \approx d_1$, $|\Delta\theta_{head}| \approx 7° \rightarrow 43° < |\phi_{az}| < 180°$
$d \approx d_2$, $|\Delta\theta_{head}| \approx 16° \rightarrow 55° < |\phi_{az}| < 180°$
$d \approx d_3$, $|\Delta\theta_{head}| \approx 24° \rightarrow 64° < |\phi_{az}| < 180°$
$d \approx d_4$, $|\Delta\theta_{head}| \approx 30° \rightarrow 67° < |\phi_{az}| < 180°$
$d_1$
$d_2$
$d_3$
$d_4$
0.2mm
D
$R^2$
Bout distance [mm]
E
### of ambiguous positions given $\Delta\theta_{head}, d$
% of total bouts
Bout distance [mm]
$\Delta\theta_{head}$ and $d$ cannot predict $\phi_{az}$ in 16% of bouts
F
$\Delta\theta_{head} \approx 0$, $\phi_{az} \approx 0$
$d \approx d_2$
$d \approx d_3$
$d \approx d_4$
Same $\Delta\theta_{head}$, same $\phi_{az}$, different $d$
$|\Delta\theta_{head}| < 5$ and $|\phi_{az}| < 5$
in 13% of bouts
$\Delta\theta_{head}$ and $\phi_{az}$ cannot predict $d$
in (at least) 13% of bouts

**Supplementary Figure 3. Bout outcomes cannot be fully represented by the change in heading angle and distance, or by the change in heading angle and azimuthal angle. A:** The relation between bout azimuthal angle and the change in heading angle was distance-dependent. Each dot represents a bout. Bouts were divided into distance groups where each group was based on 15th percentile and color-coded accordingly. The respective distance division was : 0-0.24mm, 0.24mm-0.37mm, 0.37mm-0.49mm, 0.49mm-0.65mm, 0.65mm-0.88mm, 0.88mm-1.6mm, above 1.6mm. **B:** The relation between bout azimuthal angle and change in heading angle in the first six distance groups showed a sigmoidal relation. The range of change in heading angles increased with the increase in bout distance (note the different y-axis scales). This sigmoidal relation was captured by a hyperbolic tangent (black curve) with high goodness of fit for bout distance greater than 0.2 ($R^2 = 0.55, 0.85, 0.91, 0.93, 0.95, 0.93$ for the different distance groups respectively). Dashed vertical grey bars indicate the azimuthal angles where the change in heading angle was saturated while the azimuth kept increasing. Therefore in these grey regions, the use of heading angle and distance cannot resolve the position of the fish due to varying azimuthal angles. **C:** Left: A zoom-in panel of Figure 2A from the main text, with all bouts with distances of 0.49mm-0.65mm marked between the two purple circles. Therefore, bout distance for all bouts between the two purple circles is approximately similar. Starting from the top area of the ring created by the two purple circles, as we move along the circle (to the left or right) the azimuthal angle increases in absolute value. For small azimuthal angles, the change in heading angle (color of the bout) increases with the azimuthal angle. However, as we gradually move to the right or left we enter the the saturated grey regime of the purple ring, which matches the grey regime in panel B purple rectangle. In this saturated regime, the change in heading angle is saturated and does not change for a wide range of azimuthal angles. Therefore, bout distance and change in heading angle are the same for all bouts in the grey area, and azimuthal angle cannot be inferred from them. Right: The range of the saturated areas for each distance group shown in panel C. Each distance group has an area on its respective ring, where the azimuthal angle is ambiguous given the distance and the change in heading angle. **D:** Goodness of the sigmoid fit, represented by $R^2$ for finer distance groups of 0.1 mm increments, showed that this mathematical relation accurately captured the data with $R^2 > 0.8$ for all bout distances above 0.2 mm. The fit for distances smaller than 0.2 mm was noisier because the maximal heading angle for this distance was only $7.4°$. **E:** The number of bouts with ambiguous azimuthal angle given the change in heading angle and distance decreased with distance (black curve), and the overall fraction of these bouts was 16% of all recorded bouts (cumulative distribution is shown in grey curve). **F:** Bout azimuthal angle and heading angle cannot describe the entire repertoire. For forward bouts, the azimuthal angle and change in heading angle are close to zero, irrespective of the bout distance. Therefore, in these cases bout distance cannot be inferred from the change in heading angle and the azimuthal angle. Bouts with both angles below 5 degrees constitute 13% of the data.

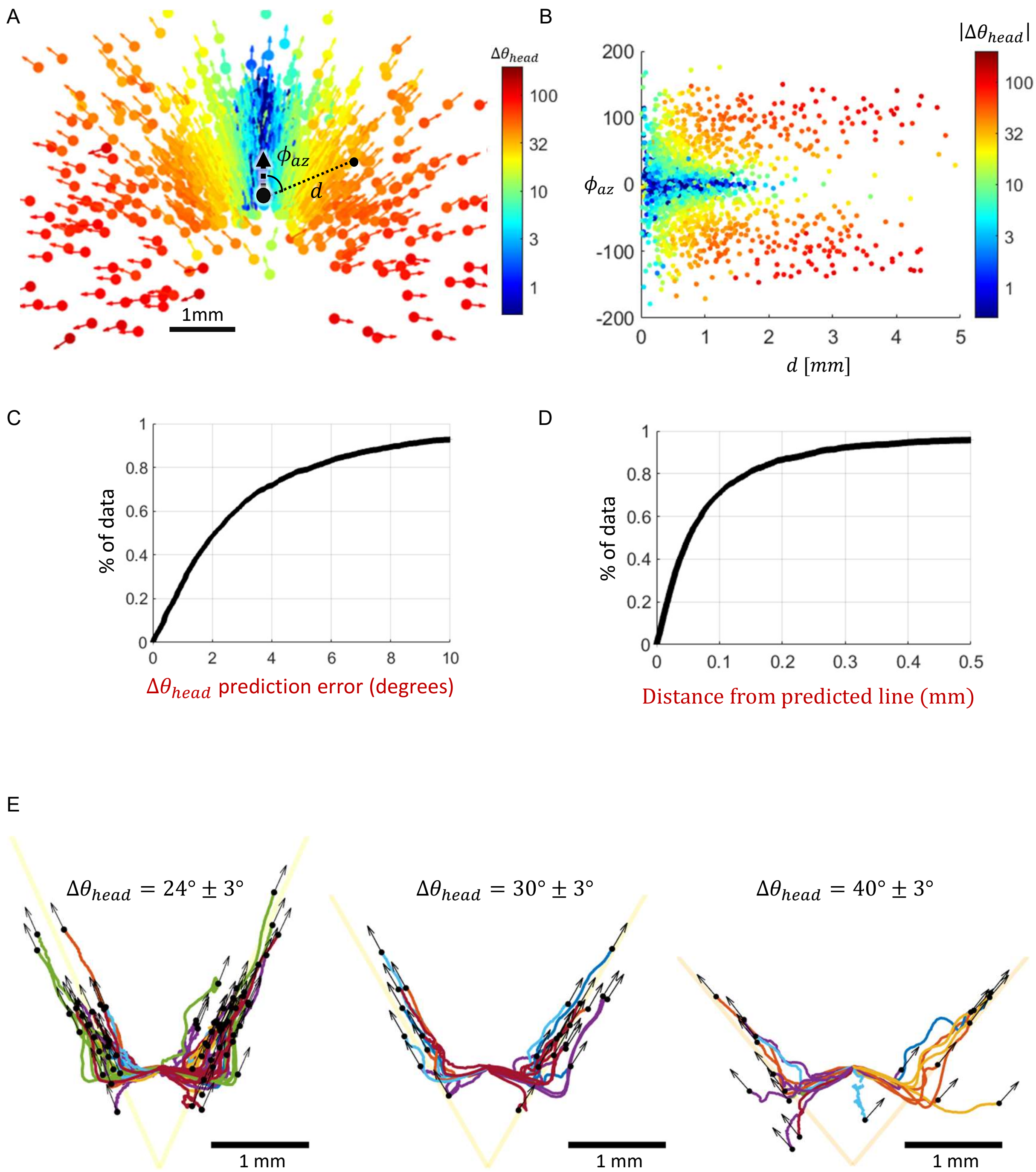


**Supplementary Figure 4. Robust model performance for older fish. A:** The repertoire of outcomes for 14-15 dpf fish is qualitatively similar to 5-7 dpf fish. Bout outcomes aligned according to the pre-bout position, similar to Figure 2A. Data contains 13 fish with 2545 bouts. **B:** The same repertoire can be represented as the change in heading angle (color) as a function of positional parameters (distance and azimuthal angle). Compare to Figure 2B. **C-D:** The model prediction of the change in heading angle given post-bout position (C) and the prediction of the linear line for each specific change in heading angle (D). Results match the performance of the model for the 5-7 dpf fish (See Figure S2A-B). 72% of the bouts show a less than $4°$ error in predicting change in heading angle given the post-bout position. 87% of bouts show a less than 0.2mm error in position given the change in heading angle. **E:** Trajectories of the point between the eyes for 14-15 dpf fish, for different changes in heading angle. Linear yellow lines are the theoretical lines predicted by the model for three example changes in heading angle ($24°$, $30°$ and $40°$). Respectively, 82%, 83% and 67% of post-bout positions were aligned along the predicted lines with an error smaller than 0.2 mm. Compare to Figure 3C, S2D.

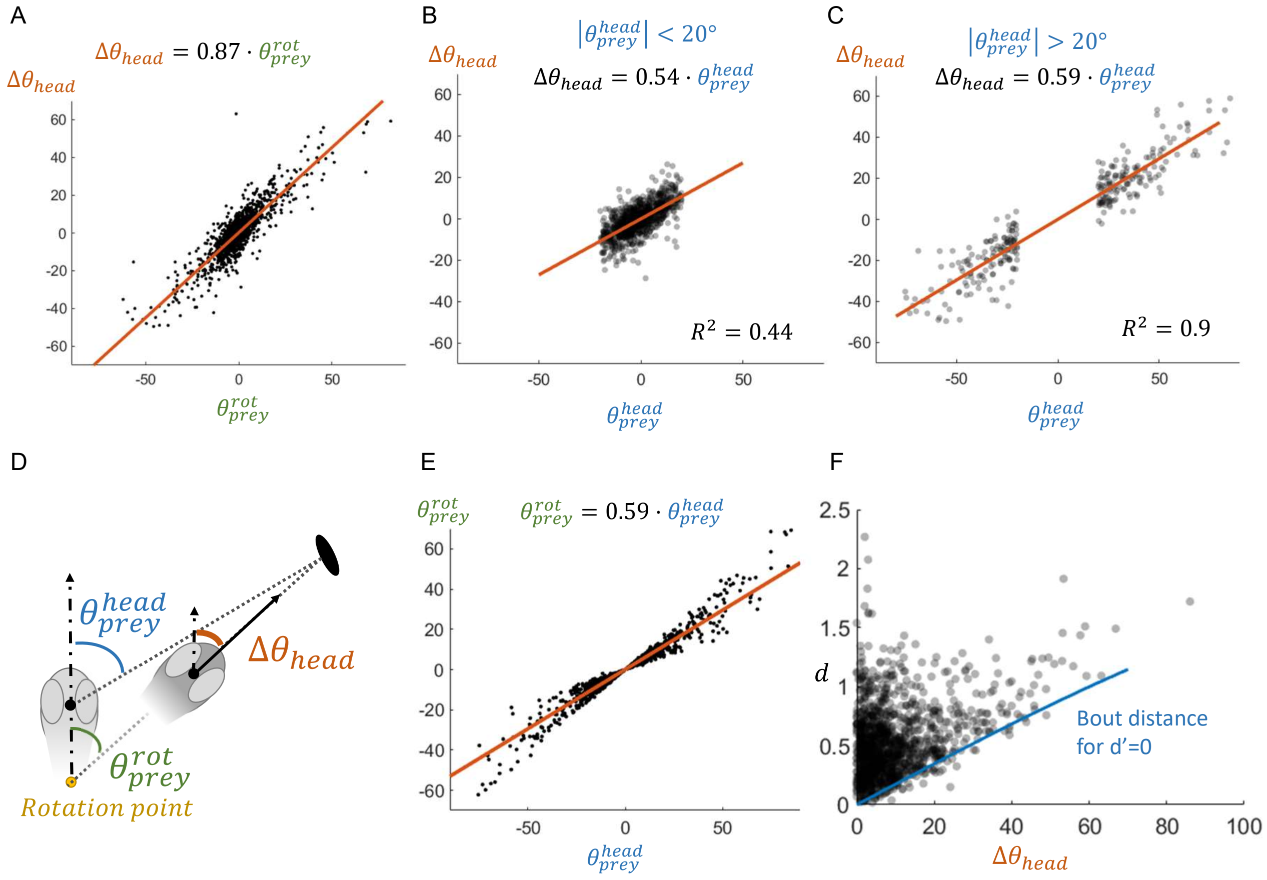


**Supplementary Figure 5. The relation between the change in heading angle and the prey angle. A:** Linear fit for change in heading angle vs prey angle relative to $y_{rot}$ ($\theta^{rot}_{prey}$), without separating the data into smaller and larger angles. **B-C:** Linear fit for change in heading angle vs prey angle relative to the point between the eyes $\theta^{head}_{prey}$ , separated into smaller and larger angles shows no difference in slope. **D:** the relation between the change in heading angle was studied relative to the prey angle with respect to the pre-bout head position or the pre-bout rotation point. **E:** Comparing the prey angle relative to the point between the eyes ($\theta^{head}_{prey}$, x-axis) to the prey angle relative to $y_{rot}$ ($\theta^{rot}_{prey}$ y-axis). The fit shows that on average the prey angle relative to $y_{rot}$ is smaller by 60%. Therefore, on average, rotating in 60% of the prey angle relative to the midpoint between the eyes ($\theta^{head}_{prey}$) is equivalent to rotating the full prey angle when measured relative to $y_{rot}$ ($\theta^{rot}_{prey}$). This suggests the fish will always be in front of the prey, regardless of the traveled distance along the theoretical line given a specific change in heading angle. **F:** Bout distance as a function of the selected change in heading angle shows that distance selection is independent of the change in heading angle. The rotation around the rotation point determines a minimal bout distance (measured relative to the point between the eyes), which increases with the change in heading angle (blue line)

A

$\theta_{head}$ $vs$ $\theta_{prey}^{rot}$ slopes for different separations of the data

| | $d_{50} = 2.34\ mm$ (50% of data) | | $d_{75} = 3.15\ mm$ (75% of data) | | $d_{90} = 4\ mm$ (90% of data) | |
|---|---|---|---|---|---|---|
| | $d_{prey} < d_{50}$ | $d_{prey} > d_{50}$ | $d_{prey} < d_{75}$ | $d_{prey} > d_{75}$ | $d_{prey} < d_{90}$ | $d_{prey} > d_{90}$ |
| $\theta_{prey}^{rot} < 5°$ (50% of data) | 1.18 ± 0.25 | 0.79 ± 0.22 | 1.07 ± 0.2 | 0.81 ± 0.33 | 1.04 ± 0.18 | 0.68 ± 0.72 |
| $\theta_{prey}^{rot} > 5°$ | 0.97 ± 0.05 | 0.82 ± 0.03 | 0.95 ± 0.04 | 0.79 ± 0.04 | 0.92 ± 0.03 | 0.75 ± 0.06 |
| $\theta_{prey}^{rot} < 10°$ (75% of data) | 1.12 ± 0.12 | 0.82 ± 0.1 | 1.03 ± 0.09 | 0.79 ± 0.14 | 1.00 ± 0.08 | 0.77 ± 0.26 |
| $\theta_{prey}^{rot} > 10°$ | 0.96 ± 0.06 | 0.82 ± 0.04 | 0.95 ± 0.04 | 0.79 ± 0.05 | 0.91 ± 0.04 | 0.75 ± 0.07 |
| $\theta_{prey}^{rot} < 20°$ (90% of data) | 1.12 ± 0.08 | 0.86 ± 0.06 | 1.05 ± 0.06 | 0.79 ± 0.08 | 1.00 ± 0.05 | 0.77 ± 0.15 |
| $\theta_{prey}^{rot} > 20°$ | 0.91 ± 0.07 | 0.81 ± 0.05 | 0.91 ± 0.06 | 0.79 ± 0.06 | 0.88 ± 0.05 | 0.75 ± 0.09 |

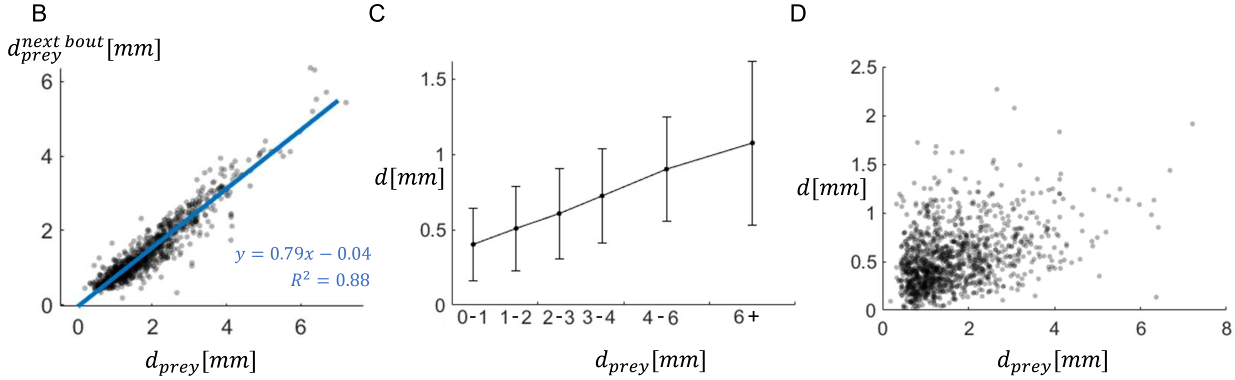


**Supplementary Figure 6. The relation between the change in heading angle and prey angle also depends on prey distance A:** Linear slopes and confidence intervals for splitting the data according to 50th, 75th and 90th percentile for both prey angle and prey distance. Middle panel ($\theta_{prey}^{rot} > 10°, d_{prey} > 3.15mm$) matches the data shown in Figure 5D. Slopes larger than 0.95 are colored in orange, and slopes smaller than 0.85 are colored in blue. In most partitions of the data, slopes are similar for smaller and larger angles but are different for smaller and larger distances. This suggests that the change in heading angle depends on the prey angle in a prey distance-dependent manner. **B:** pre- and post-bout prey distance along the event shows a decrease in the distance to the prey. **C:** On average, the selection of bout distance as a function of the prey distance shows a linear relation with a substantial variance. **D:** The selection of bout distance with respect to a given prey distance at the single bout level varies greatly between bouts.
